# The impact of non-additive genetic associations on age-related complex diseases

**DOI:** 10.1101/2020.05.12.084608

**Authors:** Marta Guindo-Martínez, Ramon Amela, Silvia Bonàs-Guarch, Montserrat Puiggròs, Cecilia Salvoro, Irene Miguel-Escalada, Caitlin E Carey, Joanne B. Cole, Sina Rüeger, Elizabeth Atkinson, Aaron Leong, Friman Sanchez, Cristian Ramon-Cortes, Jorge Ejarque, Duncan S Palmer, Mitja Kurki, FinnGen Consortium, Krishna Aragam, Jose C Florez, Rosa M. Badia, Josep M. Mercader, David Torrents

## Abstract

Genome-wide association studies (GWAS) are not fully comprehensive as current strategies typically test only the additive model, exclude the X chromosome, and use only one reference panel for genotype imputation. We implemented an extensive GWAS strategy, GUIDANCE, which improves genotype imputation by using multiple reference panels, includes the analysis of the X chromosome and non-additive models to test for association. We applied this methodology to 62,281 subjects across 22 age-related diseases and identified 94 genome-wide associated loci, including 26 previously unreported. We observed that 27.6% of the 94 loci would be missed if we only used standard imputation strategies and only tested the additive model. Among the new findings, we identified three novel low-frequency recessive variants with odds ratios larger than 4, which would need at least a three-fold larger sample size to be detected under the additive model. This study highlights the benefits of applying innovative strategies to better uncover the genetic architecture of complex diseases.

## Introduction

Genome-wide association studies (GWAS) have been successful in identifying thousands of associations between genetic variation and human complex diseases and traits ^1^. Nevertheless, for most complex diseases, only a small fraction of their genetic architecture is known and a small amount of the estimated heritability is explained ^2^. Variants that individually have small contributions to the risk of disease, and/or are rare in the population, are often missed by the majority of GWAS even though their role in the pathophysiology of complex diseases can be crucial. Some of the current limitations of GWAS could be overcome by increasing sample sizes and, as recently demonstrated, by applying more comprehensive analytical methods with improved imputation strategies ^3^. Though the increase of sample size might allow the detection of more genetic signals, it also imposes major methodological and computational requirements. These can require scientists to restrict and simplify the analysis by limiting it to autosomal chromosomes, a single reference panel for imputation, and a single (additive) inheritance model for association testing, leaving a relevant fraction of the genetic architecture of the disease unexplored ^4^.

The genetic variants that modify the risk to develop a particular complex disease may contribute to the final phenotype through different functional mechanism defined by a particular model of inheritance, which is further reflected in a characteristic distribution of affected alleles across patients and healthy individuals in GWAS. For example, the additive inheritance model, which is often the only genetic model tested, assumes that the risk of the disease is proportional to the number of risk alleles in an individual, i. e., that the effect of the heterozygous genotype is halfway between the two possible homozygous genotypes. However, some variants follow non-additive inheritance models, which include dominant, recessive and heterodominant. The additive model is expected to capture a large fraction of the genetic risk for disease ^5^ and can identify some variants that follow non-additive inheritance patterns. However, the additive model is not sufficient to provide a comprehensive overview of the genetic architecture of diseases. In particular, most GWAS may have insufficient power to identify low-frequency variants that show recessive effects ^6, 7^. The importance of evaluating non-additive inheritance models is well reported in the context of Mendelian diseases ^8^ and occasionally for complex traits as well, such as the recessive effects of the *FTO* locus in obesity ^9^, the *ITGA1* ^10^, *TBC1D4* ^11^ and *CDKAL1* ^9, 12^ genes in type 2 diabetes, as well as the known non-additive effects of HLA haplotypes in autoimmune diseases ^13^ and ulcerative colitis ^14^. The increasing ability to capture low-frequency variants using modern imputation reference panels and the need to uncover the still missing heritability estimated for most complex diseases, call for comprehensive association strategies that should include, among other improvements, the analysis of non-additive inheritance models.

To fill this gap and to determine the prevalence and contribution of the different inheritance patterns involved in the genetic architecture of complex diseases, we have designed and implemented a comprehensive strategy for genetic association analysis that combines dense imputation from multiple reference panels with association testing under five different inheritance models across multiple phenotypes. We have applied this strategy to the Kaiser Permanente Research Program on Genes, Environment and Health: A Genetic Epidemiology Research on Adult Health and Aging (GERA) cohort ^15^, which includes 62,281 subjects from European ancestry and 22 diseases.

Finally, we release here both the summary statistics for all the models of inheritance as well as the complete methodology, provided to the community as an easy-to-use and standalone pipeline. This pipeline allows for analysis of existing and newly generated GWAS data with better efficiency and more comprehensive testing, improving the chances of variant discovery.

## Results

In order to assess the potential benefits of applying more in-depth GWAS methodologies to available genetic datasets, and to investigate the relative contribution of different inheritance models to the risk to develop complex diseases, we have applied a global analysis strategy to the GERA cohort, an age-related disease-based cohort with an average age of 63, well powered to study a broad range of clinically defined age-related conditions. By using this particular cohort, we expect to minimize a possible loss of power due to the misclassification of controls, as often happens in datasets with younger individuals that can include cases at pre-disease stages classified as controls.

### Genotype Imputation and association testing using multiple reference panels

After applying strict genetic quality control to the GERA cohort (see Methods), we retained 56,637 individuals with European ancestry for further downstream analysis (Supplementary Table 1). To cover the maximum number and type of genetic variants, we next applied an extensive imputation strategy with four reference panels: the Genome of the Netherlands (GoNL) ^16, 17^, the UK10K Project ^18^, the 1000 Genomes Project (1000G) phase 3 ^19^ and Haplotype Reference Consortium (HRC) ^20^, and imputed 11.2 M, 11.4 M, 13.1 M, and 11.7 M high quality imputed variants (IMPUTE2 ^21^ info score ≥ 0.7 and minor allele frequency [MAF] ≥ 0.001) with each panel, respectively. After combining the results of the four reference panels by choosing, for each variant, the panel that provided the highest imputation accuracy, we retained a total of 16,059,686 variants covering all the autosomes and the X chromosome (Figure 1a). This strategy was particularly powerful to impute 2.6 M and 5.5 M high quality, low-frequency (0.05 > MAF > 0.01) and rare variants (0.01 > MAF > 0.001), respectively, as well as 1.6 M indels. Note that as many as 684,393 common variants (MAF ≥ 0.05), 255,106 low-frequency, 1.7 M rare, and all indels (1.6 M) would be missed if only the HRC reference panel was used. This highlights the benefit of combining different reference panels for comprehensive association testing (Figure 1b).

**Table 1.**
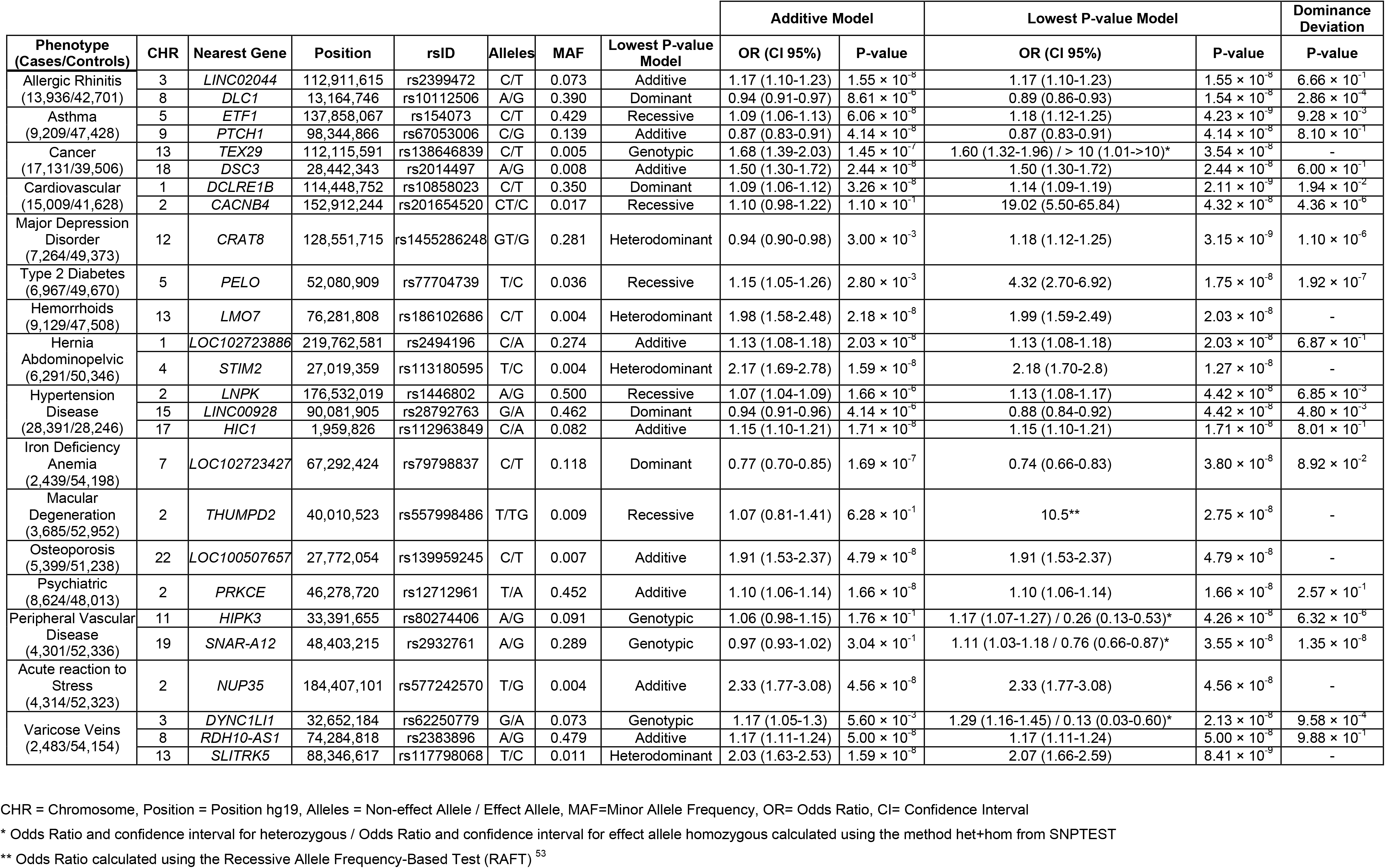
New associations from the GERA cohort analysis.

**Figure 1.**
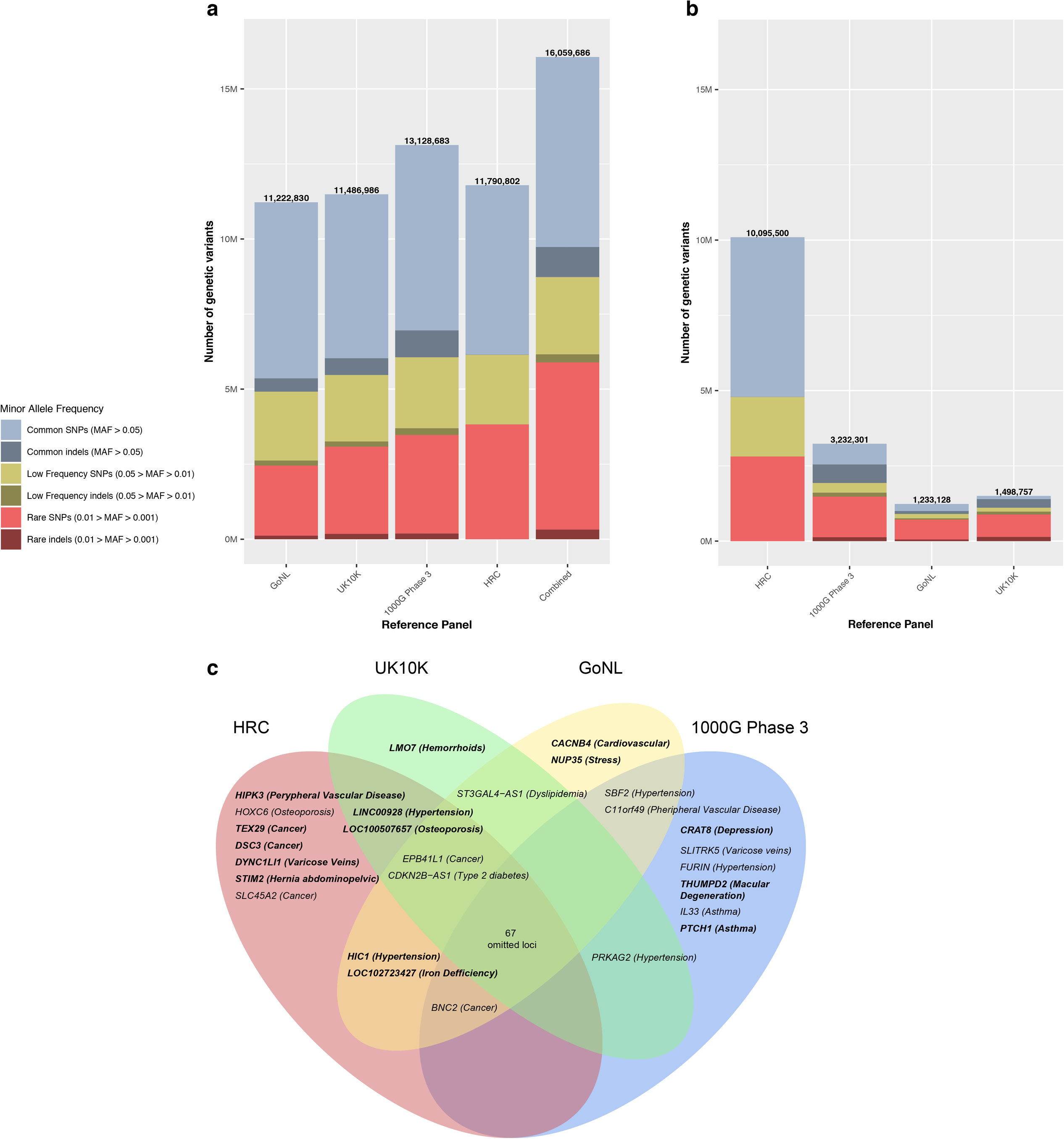
Graphical representation illustrating the benefits of combining the results from different reference panels. **a** Comparison of the number of variants after the imputation with four reference panels (info score ≥ 0.7), and combining them, colored according to MAF and varianat type (SNP vs alternative forms of variation, such as indels). As shown in the bar plot, combining the results from the four reference panels increased the final set of variants for association testing when compared with the results for each of the panels alone (GoNL, UK10K, 1000G Phase 3 or HRC), especially in the low and rare frequency spectrum. For example, we covered up to 5.5 M rare variants (0.01 > MAF > 0.001) by combining panels, while only 2,3 M, 2,9 M, 3,2 M and 3,8 M of rare variants were imputed independently with GoNL, UK10K, 1000G phase 3 and HRC, respectively. **b** Comparison of the contribution of each reference panel in the combined results. Each bar represents the number of variants that had the best imputation accuracy for a given reference panel. As shown in the figure, although the HRC panel showed overall higher imputation scores, as it provided around 10 of the final 16 M variants, the contribution of the other reference panels, primarily with non-SNP variants, was substantial. Indels seen in the bar plot for HRC correspond to genotyped indels. All variants with info score < 0.7, MAF < 0.001 and HWE for controls *p* < 1.0 × 10^−6^ were filtered. **c** Venn Diagram illustrating the loci that identified by each reference panel. New loci are depicted in bold. As shown in this figure, only 67 of the 94 GWAS significant loci were identified by all four reference panels, while 27 of them (28.7%) were only identified by one, two or three of the four panels.

We next tested all the 16 M variants for association with the 22 conditions available in the GERA cohort considering the entire genome and five different inheritance models (Supplementary Figure 1-22). This analysis identified 94 independent loci associated with 17 phenotypes at a genome-wide significance level (*p* < 5.0 × 10^−8^) of which 63 for 14 phenotypes were also experiment-wide significant (*p* < 2.0 × 10^−8^) after considering correction for the different models of inheritance (see methods) (Supplementary Table 2). According to the GWAS catalog, 68 of the 94 genome-wide significant loci had been previously reported to be associated with the same disease (Supplementary Table 3), whereas 26 of them correspond to previously unreported loci with associations across 16 phenotypes (Table 1). Of these new loci, 16 correspond to common, 3 to low-frequency, and 7 to rare variants. Interestingly, only a fraction of the 26 new loci would have been genome-wide significant by using individual imputation panels (Figure 1c), namely 20/26 using HRC, 14/26 using 1000G Phase 3, 14/26 using UK10K or 15/26 using GoNL. In addition, the lead marker for three of the novel signals is an indel, further confirming the benefits of combining multiple panels with our approach.

**Table 2.** Replication of new associations with UK Biobank.

|  |  |  | Stage 1. Discovery |  |  |  | Stage 2. Replication |  |  |  | Stage 1 + Stage 2. Meta-analysis |  |  |  |  |  |
| --- | --- | --- | --- | --- | --- | --- | --- | --- | --- | --- | --- | --- | --- | --- | --- | --- |
| CHR | rsID (Alleles)<br>(MAF) | Best Model | Phenotype<br>(Cases/Controls) | Additive |  | Best Model |  | Field<br>(Cases/Controls or<br>Sample Size) | Additive |  | Lowest p-value model |  | Additive |  | Lowest p-value model |  |
|  |  |  |  | OR (CI 95%) | P-value | OR (CI 95%) | P-value |  | OR (CI 95%) | P-value | OR (CI 95%) | P-value | OR (CI 95%) | P-value | OR (CI 95%) | P-value |
| 18 | rs2014497<br>(A/G) (0.008) | Additive | Cancer<br>(17,131/39,506) | 1.50 (1.30-1.72) | 2.44×10<sup>-8</sup> | 1.50 (1.30-1.72) | 2.44×10<sup>-8</sup> | Self-reported:<br>chronic lymphocytic<br>(237/360,904) | 2.13 (1.14-3.97) | 3.50×10<sup>-2</sup> | 2.13 (1.14-3.97) | 3.50×10<sup>-2</sup> | 1.52 (1.33-1.74) | 1.60×10<sup>-9</sup> | 1.52 (1.33-1.74) | 1.60×10<sup>-9</sup> |
|  |  |  |  |  |  |  |  | Self-reported:<br>kidney/renal cell<br>cancer<br>(473/360,668) | 1.75 (1.07-2.86) | 4.25×10<sup>-2</sup> | 1.75 (1.07-2.86) | 4.25×10<sup>-2</sup> | 1.51 (1.32-1.73) | 1.49×10<sup>-9</sup> | 1.51 (1.32-1.73) | 1.49×10<sup>-9</sup> |
|  |  |  |  |  |  |  |  | C69 Malignant<br>neoplasm of eye and<br>adnexa<br>(146/361,048) | 2.51 (1.19-5.3) | 3.56×10<sup>-2</sup> | 2.51 (1.19-5.3) | 3.56×10<sup>-2</sup> | 1.52 (1.33-1.75) | 1.95×10<sup>-9</sup> | 1.52 (1.33-1.75) | 1.95×10<sup>-9</sup> |
| 1 | rs2494196<br>(C/A) (0.274) | Additive | Hernia<br>Abdominopelvic<br>(6,291/50,346) | 1.13 (1.08-1.18) | 2.03×10<sup>-8</sup> | 1.13 (1.08-1.18) | 2.03×10<sup>-8</sup> | Self-reported:<br>umbilical hernia<br>(328/360,813) | 1.42 (1.21-1.67) | 2.31×10<sup>-5</sup> | 1.42 (1.21-1.67) | 2.31×10<sup>-5</sup> | 1.15 (1.10-1.19) | 5.35×10<sup>-11</sup> | 1.15 (1.10-1.19) | 5.35×10<sup>-11</sup> |
|  |  |  |  |  |  |  |  | K40 Inguinal hernia<br>(13,365/347,829) | 1.09 (1.06-1.12) | 3.95×10<sup>-10</sup> | 1.09 (1.06-1.12) | 3.95×10<sup>-10</sup> | 1.10 (1.08-1.12) | 7.78×10<sup>-17</sup> | 1.10 (1.08-1.12) | 7.78×10<sup>-17</sup> |
|  |  |  |  |  |  |  |  | K41 Femoral hernia<br>(475/360,719) | 1.44 (1.26-1.64) | 1.24×10<sup>-7</sup> | 1.44 (1.26-1.64) | 1.24×10<sup>-7</sup> | 1.16 (1.11-1.21) | 2.26×10<sup>-12</sup> | 1.16 (1.11-1.21) | 2.26×10<sup>-12</sup> |
|  |  |  |  |  |  |  |  | K42 Umbilical hernia<br>(2,623/358,571) | 1.29 (1.22-1.37) | 1.14×10<sup>-17</sup> | 1.29 (1.22-1.37) | 1.14×10<sup>-17</sup> | 1.19 (1.15-1.22) | 2.94×10<sup>-22</sup> | 1.19 (1.15-1.22) | 2.94×10<sup>-22</sup> |
|  |  |  |  |  |  |  |  | K43 Ventral hernia<br>(2,470/358,724) | 1.18 (1.11-1.25) | 1.77×10<sup>-7</sup> | 1.18 (1.11-1.25) | 1.77×10<sup>-7</sup> | 1.15 (1.11-1.19) | 1.99×10<sup>-14</sup> | 1.15 (1.11-1.19) | 1.99×10<sup>-14</sup> |
| 2 | rs557998486<br>(T/TG) (0.009) | Recessive | Macular<br>Degeneration<br>(3,685/52,952) | 1.07 (0.81-1.41) | 6.28×10<sup>-1</sup> | 10.5* | 2.75×10<sup>-8</sup> | Eye<br>problems/disorders:<br>Macular<br>degeneration<br>(2,726/115,164) | 0.98 (0.72-1.32) | 8.81×10<sup>-1</sup> | 7.58 (1.54-37.32) | 4.1×10<sup>-2</sup> | 1.01(0.82-1.24)** | 7.91×10<sup>-1***</sup> | 26.51(7.57-92.85)** | 3.29×10<sup>-8***</sup> |
| 5 | rs77704739<br>(T/C) (0.036) | Recessive | Type 2 Diabetes<br>(6,967/49,670) | 1.15 (1.05-1.26) | 2.80×10<sup>-3</sup> | 4.32 (2.70-6.92) | 1.75×10<sup>-8</sup> | Self-reported:<br>diabetes<br>(14,114/347,027) | 1.03 (0.97-1.09) | 3.87×10<sup>-1</sup> | 1.88 (1.35-2.6) | 4.95×10<sup>-4</sup> | 1.06 (1.01-1.12) | 1.78×10<sup>-2</sup> | 2.46 (1.88-3.21) | 4.68×10<sup>-11</sup> |

CHR = Chromosome, Position = Position hg19, Alleles = Non-effect Allele / Effect Allele, MAF= Minor Allele Frequency, OR= Odds Ratio
\* Odds Ratio calculated using the Recessive Allele Frequency-Based Test (RAFT)
\*\* Obtained through a mega-analysis with UK Biobank using the "expected" method from SNPTTEST
\*\*\* Obtained using METAL method "SAMPLESIZE" to combine the p-values taking into account the sample size and direction of effect.

### Identification of recessive variants with large effects

The implementation of refined GWAS strategies not only increases the number of associated variants, but also allows the identification of loci with large impact on the disease. Among the variants that were not detected under the additive model, and hence are expected to be missed by the majority of current GWAS, we highlight three variants with remarkably large recessive effects. First, an intronic indel in the *CACNB4* gene, rs201654520, associated with a nearly twenty-fold increase in risk for cardiovascular disease (MAF= 0.017, OR [CI 95%] = 19.0 [5.5 - 65.8], *p* = 4.3 × 10^−8^). *CACNB4* encodes the β4 subunit of the voltage-dependent calcium channel. This subunit contributes to the flux of calcium ions into the cell by increasing peak calcium current and triggering muscle contraction. Interestingly, an intronic single nucleotide polimorphism (SNP) within *CACNB4*, rs150793926, was associated with idiopathic dilated cardiomyopathy in African Americans ^22^, but this variant is not in linkage disequilibrium (LD) with rs201654520 (LD r^2^ ^23^ = 0.0016).

A second recessive variant with large effect, rs77704739, near the *PELO* gene, is associated with a four-fold risk for type 2 diabetes (MAF= 0.036, OR [CI 95%] = 4.3 [2.7 - 6.9], *p* = 1.75 × 10^−8^). We also found this variant associated with type 2 diabetes (OR-recessive [95% CI] = 1.9 [1.4 - 2.6], p = 4.95 × 10^−4^) and metformin use (OR-recessive [95% CI] = 2.3 [1.6 - 3.4], p = 3.8 × 10^−5^) in the UK Biobank ^24^ (Supplementary Table 4), also only under the recessive model. An independent signal that is about 112 K base pairs away (rs870992, LD r^2^ = 0.0009) was previously associated with type 2 diabetes in the Greenlandic population, also with a recessive effect ^10^. To provide insights into the underlying molecular mechanisms in disease, we interrogated comprehensive catalogues of genetic effects on gene expression; eQTLGen Consortium ^25^ and GTEx ^26^. The rs77704739 variant was significantly associated with gene expression of *PELO* in multiple tissues, including diabetes-relevant tissues such as adipose tissue, skeletal muscle, and pancreas. Colocalization analyses showed a probability higher than 0.8 in several tissues, including subcutaneous adipose tissue and skeletal muscle, suggesting this gene as the effector transcript (Figure 3a, 3b, and Supplementary Table 5). In addition, we found that the lead variants in the *PELO* locus overlap with active promoter annotations in human pancreatic islets and open chromatin sites highly-bounded by islet specific transcription factors ^27, 28^ (Figure 3c).

Third, a rare indel, rs557998486, located near the *THUMPD2* gene, is associated with age-related macular degeneration (MAF= 0.009, OR = 10.5, *p* = 2.75 × 10^−8^). Also under the recessive model in UK Biobank, this indel was associated with age-related macular degeneration (OR [CI 95%] = 7.6 [1.5-37.3], *p* = 4.1 × 10^−2^), eye surgery (beta [CI 95%] = 1.6 [0.6-2.6], *p* = 1.17 × 10^−3^) (Supplementary Table 4), and C-reactive protein, a known biomarker for macular degeneration ^29^ (beta [CI 95%] =1.1 [0.7 - 1.5], *p* = 1.15 × 10^−4^) (Supplementary Table 6). Interestingly, the fact that we found no SNPs in LD with this lead indel further confirms the benefits of multiple reference panel imputation strategies that include alternative forms of variation. The lead indel rs557998486 overlaps DNAse I hypersensitivity sites in retinal and iris cell lines ^30^, highlighting a candidate open chromatin region that is also predicted to be an enhancer assigned to the *THUMPD2* gene according to GeneHancer ^31^. One of the variants with the highest LD with rs557998486 (rs116649730, LD r^2^= 0.32) is associated with reduced expression of its nearest gene, *THUMPD2* (Z-score = −4.85, *p* = 1.25 × 10^−6^), according to eQTLGen Consortium data.

### Replication using UK Biobank and FinnGen

We sought replication of previously unreported loci using UK Biobank, a prospective cohort of ~500 K individuals aged between 40 to 69 ^24^. Given the high heterogeneity in phenotype definitions in UK Biobank compared to GERA, we tested for replication with the same phenotype and related traits (Supplementary Table 4). Compared to GERA, some of the conditions may not be ascertained or have an age at onset later than the average age at ascertainment in UK Biobank (56.52 years ^32^) which could affect the replication success. Despite these limitations, we tested the novel variants using the corresponding inheritance model, and replicated 4 new loci with the same phenotype (Table 2).

We further sought replication of the association within the *CACNB4* gene with cardiovascular disease in FinnGen, a cohort of ~218 K Finnish individuals with an average age of 63, as it includes individuals with a higher average age (63 vs 56 in UK Biobank) and the risk of developing a cardiovascular disease is well-known to increase with age ^33^. In addition, FinnGen has a precise and richer classification of this particular phenotype than UK Biobank. In brief, we tested rs201654520 for association with 47 cardiovascular endpoints. Of all the conditions tested, four (hypertensive heart disease, hypertensive heart and/or renal disease, heart failure, and right bundle-branch block) were nominally associated (*p* < 0.05). All the associations had a direction of effect consistent with the effect observed in the GERA cohort (Supplementary Figure 23a). Although there is a high heterogeneity in the phenotype definitions between cohorts, we meta-analyzed the results from these endpoints from FinnGen with the result from “cardiovascular disease” phenotype from GERA, but none of them reach the genome-wide significance (see Methods) (Supplementary Figure 23). We did not include UK Biobank in this meta-analysis as the equivalent phenotypes were not available or had less than 350 cases in UK Biobank, therefore, underpowered for a recessive analysis. Notably, when analyzing the association of rs201654520 with related quantitative traits we found that those who were homozygous for the high-risk allele had lower systolic blood pressure (p = 4.1 × 10^−3^, beta = −0.23) (Supplementary Table 4). While lower systolic blood pressure has been associated with increased risk of myocardial infarction in particular circumstances, this is not the typical direction of association, and therefore merits additional study ^34^.

We also sought replication of the recessive association of rs557998486 near *THUMPD2* gene with macular degeneration in FinnGen. While rs557998486 was associated with increased risk of macular degeneration in UK Biobank under the recessive model (OR [CI 95%] = 7.6 [1.5-37.3], p = 4.1 × 10^−2^), it was not significantly associated in the FinnGen biobank although it showed the same direction of effect. However, the meta-analysis did not reach the genome-wide significance (rs557998486 *p* = 9.6 × 10^−6^) and had a high heterogeneity (heterogeneity I^2^ = 87.1, heterogeneity *p* = 4.3 × 10^−4^).

### Detection ranges of the different inheritance models

Our findings provide an empirical overview of the detection range of five different inheritance models, and show how each of them captures a fraction of the genetic variants associated with complex traits. As indicative of the power of current genetic studies that usually only consider additive allelic effects, we found three different scenarios. Among all the 94 associated loci identified, 12 showed genome-wide significance only under the additive model, 62 under both additive and non-additive models, and 20 showed genome-wide significance only when non-additive tests were applied (Figure 2a). To further classify these variants, we tested whether any of the 62 variants associated with both additive and non-additive models deviate from additivity through a dominance deviation test ^9^. Eleven of these 62 variants (17.7%) showed significant deviation from additivity (dominance deviation test *p* < 0.05). Altogether, the dominance deviation test over the 93 autosomal loci identified 62 additive (66%) and 24 non-additive associations (25.5%) and 8 undetermined. Based on the smallest GWAS *p*-value, we further classified non-additive associations into 9 recessive, 13 dominant, 8 heterodominat and 7 genotypic (Supplementary Table 2).

**Figure 2.**
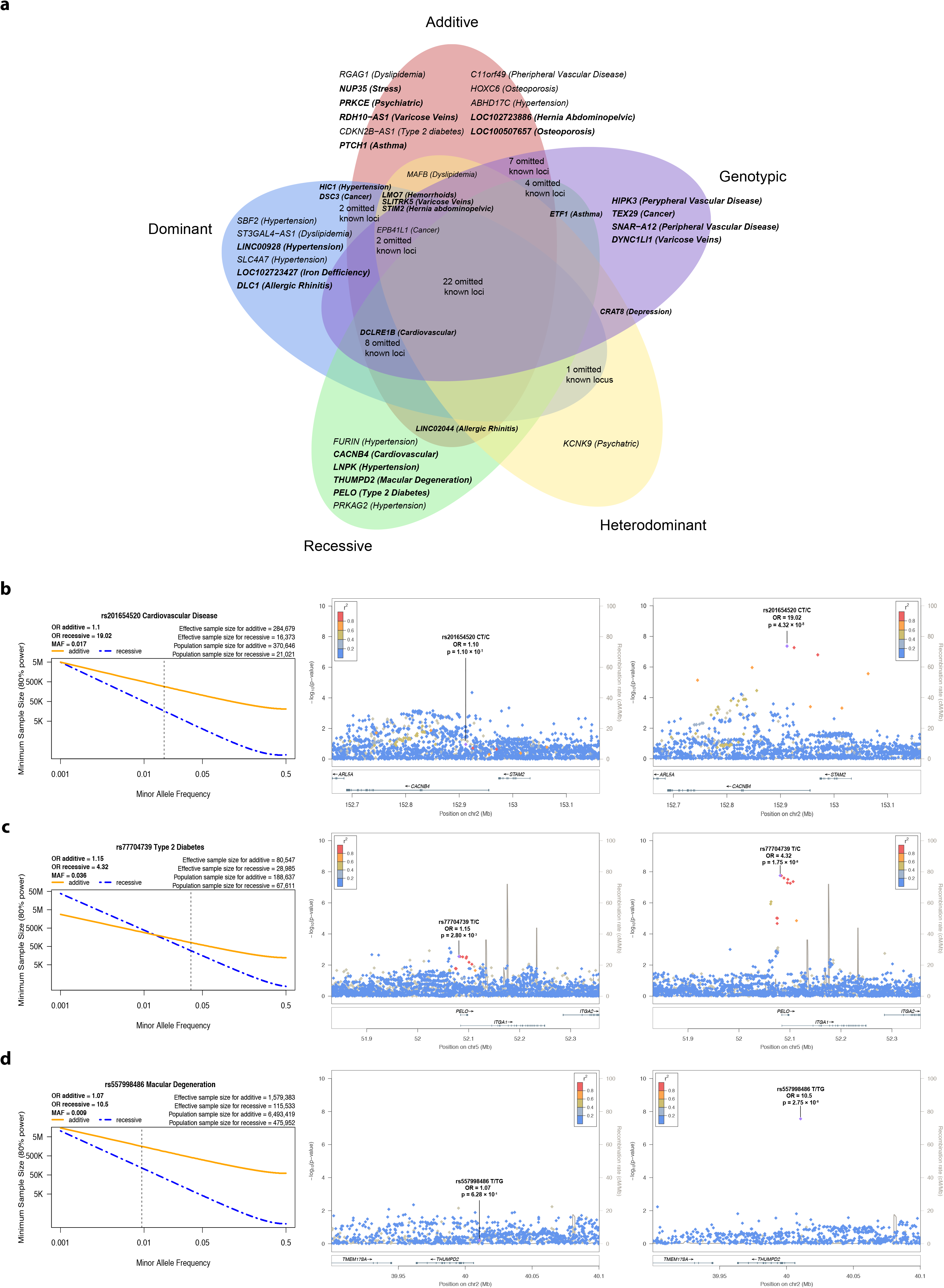
Results from the analysis of additive and non-additive inheritance models. **a** The Venn Diagram shows the number of loci that were identified when analyzing multiple inheritance models. As seen in the Venn Diagram, the strongest association for 37 of the 94 associated loci was non-additive. Moreover, the analysis of non-additive models was crucial for the identification of 14 novel (in bold) associated loci. **b** Power calculation of the rs201654520 indel in *CACNB4* associated with cardiovascular disease. The results show that the additive-based test would require a population sample size of 370,646 individuals to find this recessive association, while the population sample size needed for the recessive model was 21,021. **c** Power calculation of the rs77704739 variant near the *PELO* gene associated with type 2 diabetes. The results show that the additive-based test would require a population sample size of 188,637 individuals to find this recessive association, while the population sample size needed for the recessive model is 67,611. **d** Power calculation of the rs557998486 indel near the *THUMPD2* gene associated with age-related macular degeneration. The results show that the additive-based test would require a population sample size of 6,493,419 individuals to find this recessive association, while the population sample size for the recessive model is 157,450.

**Figure 3.**
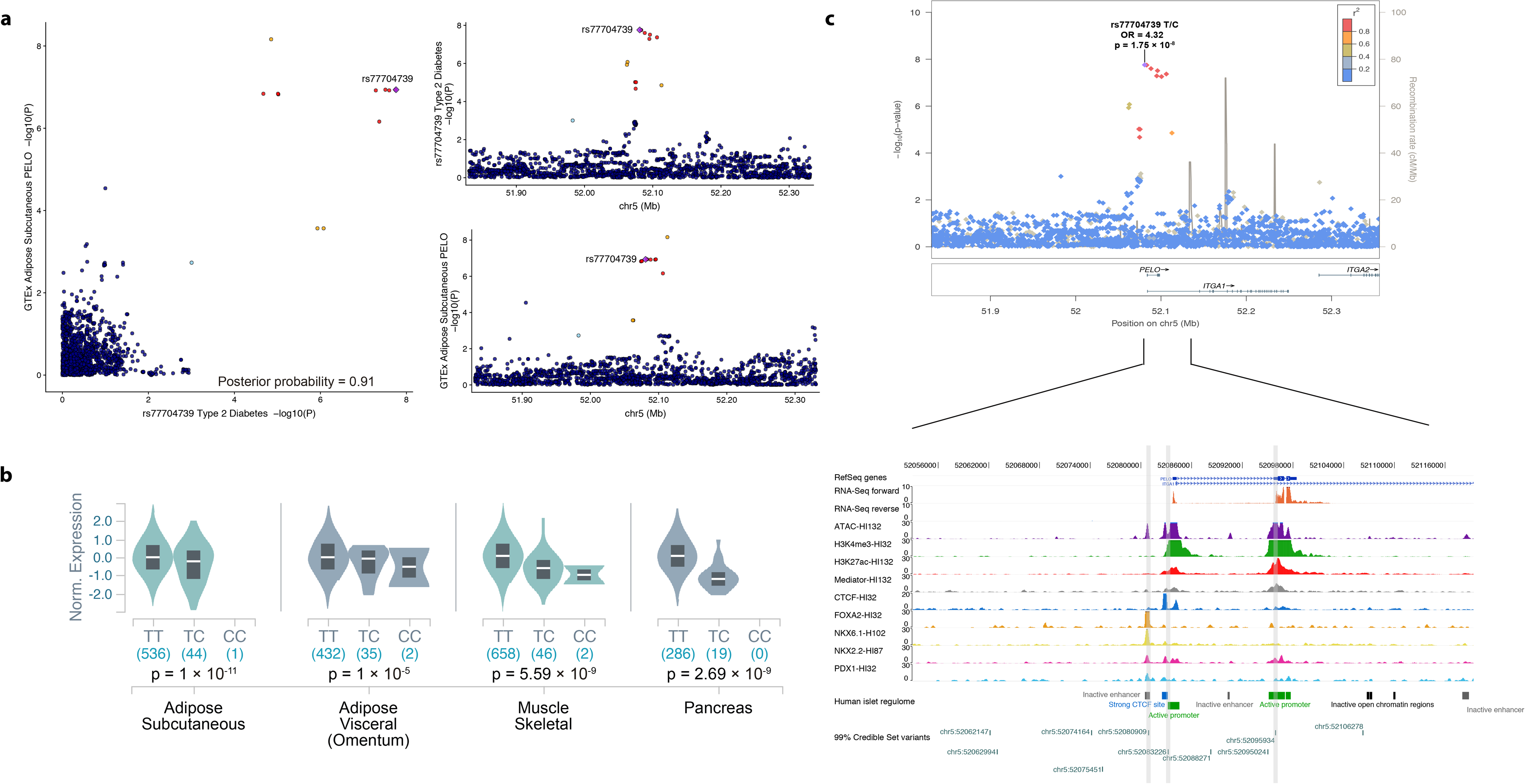
Functional characterization of the rs77704739 recessive association near the *PELO* gene. **a** Signal plot for chromosome 5 region surrounding rs77704739. Each point represents a variant, with its *p*-value from the discovery stage on a −log10 scale in the y axis. The x axis represents the genomic position (hg19). Three credible set variants are located in open chromatin sites in human pancreatic islets, one of them classified as an active promoter and one highly bounded by pancreatic islet specific transcription factors, such as PDX1, NKX2.2, NKX6.1 and FOXA2. **b** Colocalization plots from LocusCompare for the rs77704739 variant in adipose subcutaneous tissue. As seen in the plots, the signals from both eQTL data and the recessive T2D association results colocalize. c Violin plot from GTEx showing that the recessive rs77704739 variant significantly modifies the expression of *PELO* gene in subcutaneous and visceral adipose tissue, skeletal muscle and pancreas. GTEx V7 was used for colocalization analyses, whereas GTEx V8 was used to generate the violin plots.

We also found that each of the available models for association testing has a different range of detection. To identify the 94 genome-wide associated loci, the additive test, as expected, was the most sensitive model (74 loci), followed by the genotypic (59 loci), the dominant (56 loci), the recessive (43 loci) and the heterodominant (32 loci). When considering known loci, 48 of the 68 previously reported loci were identified by more than one model in our analysis, and almost half of these (22 loci) with all five models. In contrast, of the 26 newly discovered variants, only 8 were identified with multiple models, whereas the majority of them (18 loci), were detected only with the additive (6 loci), the genotypic (4 loci), the recessive (4 loci) and the dominant (3 loci) model. Of note, 13 out of 26 (50%) novel loci were only identified by non-additive models.

To further investigate to what extent the additive model captures non-additive signals, and how much this depends on sample size, we carried out power calculations on loci that were detected here only under a non-additive model, such as rs201654520 within *CACNB4* gene and rs77704739 near the *PELO* gene. These power calculations showed that the additive test would require a population sample size of at least 370,646 individuals to detect the recessive association of rs201654520 in *CACNB4* (Figure 2b), and at least 188,637 individuals to capture the recessive signal of rs77704739 near the *PELO* gene (Figure 2c), while the population sample size required for the recessive model was only 21,021 and 67,611, respectively. In this study, we were able to identify both associations with a modest sample size by using the most well-suited disease model.

### The GUIDANCE framework

We developed an integrated framework including our methodology used to analyze the GERA cohort, called GUIDANCE. GUIDANCE allows the analysis of genome-wide genotyped data in a single execution in distributed computing infrastructures without the need for extensive computational expertise or constant user intervention. The GUIDANCE workflow requires quality-controlled genotyped data as an input and provides association results, graphical outputs and statistical summaries. Integrating state-of-the-art tools with in-house code written in Java, bash and R ^35^, GUIDANCE efficiently performs large-scale GWAS, including 1) the pre-phasing of haplotypes, 2) the imputation of genotypes using multiple reference panels, 3) the association testing for different inheritance models and integrating results from different panels, 4) a cross-phenotype analysis when more than one phenotype is available in the cohort (Supplementary Table 7), and finally, 5) the generation of summary statistics tables and graphic representations of the results (Supplementary Figure 24), for both the autosomes and the X chromosome. While GUIDANCE can be executed as a standalone compact program it can also be used in modules (Supplementary Figure 25), which makes GUIDANCE adaptable to existing frameworks and provides an even higher level of control to users.

GUIDANCE runs in distributed computing platforms, including the cloud, without requiring a broad background in distributed environments. This is feasible since GUIDANCE was implemented on top of the COMP Superscalar Programming Framework (COMPSs) ^36^. With COMPSs, the GUIDANCE workflow was implemented as a sequential Java program containing the calls to the GWAS tools, encapsulated in Java methods, and selected as tasks, while COMPSs controls the execution of those tasks on the underlying distributed infrastructure. The source code, the pre-compiled binaries and documentation to use GUIDANCE are available at http://cg.bsc.es/guidance.

## Discussion

The increasingly large sample sizes in GWAS improve the statistical power to identify genetic variants associated with complex diseases. At the same time, the emergence of larger and denser reference panels allows genotype imputation at lower ranges of allele frequencies previously unexplored. In this study, we demonstrate the value of applying a comprehensive GWAS including denser imputation strategies, the X chromosome and non-additive association tests to an existing large-scale genetic resource, the GERA cohort. We show that by applying more powerful imputation protocols we increased the number and the type of variants tested for association, including low-frequency and rare SNPs as well as alternative forms of variation, such as indels. Our analysis in the GERA cohort shows that between 13 and 20 of the genome-wide significant associations (14-21%) would not have been identified when using a single reference panel. Likewise, our analysis in the GERA cohort demonstrates that 21% of the associations would be missed by only testing the additive model. Overall, 27.6% of associations would not have been identified by applying the commonly used HRC and additive model association testing.

We here show the potential of identifying very large effect recessive associations by maximizing the use of current reference panels and testing different inheritance models, as exemplified by the associations with type 2 diabetes, cardiovascular disease and macular degeneration with variants near *PELO*, *CACNB4*, and *THUMPD2*, respectively. This strategy opens new avenues for future analyses in large scale biobanks, as demonstrated with our power calculations, which show that even the largest available GWAS meta-analyses or biobanks would not have enough power to identify these associations using only the additive model. For example, the *CACNB4* gene, associated with cardiovascular disease, would require a sample size equivalent to 370,000 individuals when using the additive test, 17 times larger than the required sample size under a recessive analysis. After considering all the supporting evidence illustrated with many examples in this study, the results suggest that this new associations deserve future validations and follow-up analysis, and demonstrate the importance of a comprehensive analysis including non-additive models when performing GWAS.

The inclusion of non-additive associations can also have an impact on the construction of polygenic risk scores. Current polygenic scores (PRS) are calculated summing risk alleles weighted by effect sizes from GWAS results, which have typically tested only the additive model in the association test. Hence, large-scale genome-wide association data accounting for different models of inheritance and including both SNPs and alternative forms of variation, such as indels, will also be essential to develop more accurate genome-wide PRS, which would weight each of the genotype carriers appropriately, rather than weighting the heterozygous half-way between the homozygous of the effect and alternate alleles.

To easily apply this strategy to genetic studies we present GUIDANCE, a standalone and easy-to-use application that allows an efficient and comprehensive GWAS analysis in different computing platforms, such as cloud and high-performance computing architectures. In a moment where the community is facing computational and methodological challenges due to the growing complexity and size of genetic datasets, the availability of robust and complete analysis platforms can improve the efficiency of genetic studies, standardizing analysis strategies among large meta-analysis cohorts to ensure consistency.

Finally, to share our results with the community and to promote the analysis of non-additive inheritance models in GWAS, a public searchable database including additive and non-additive summary statistics for 16 M of variants and 22 phenotypes is available at the Type 2 Diabetes Knowledge Portal (http://www.type2diabetesgenetics.org and full summary statistics at http://cg.bsc.es/guidance).

## Online Methods

### GUIDANCE Workflow Description

By combining and integrating state-of-the-art GWAS analysis tools into the COMP Superscalar programming Framework (COMPSs), we developed GUIDANCE, a standalone application that performs haplotype phasing, genome-wide imputation, association testing and PheWAS analysis of large GWAS datasets (Supplementary Figure 24).

As shown in Supplementary Figure 24, GUIDANCE’s workflow starts with quality-controlled genotype data and ends with providing association results, graphical outputs and statistical summaries.

Once everything is settled in the GUIDANCE configuration file, GUIDANCE performs an efficent two-stage imputation procedure, by pre-phasing the genotypes into whole haplotypes followed by genotype imputation itself ^21^. SHAPEIT2 ^37^ or EAGLE2 ^38^ and IMPUTE2 ^39^ or MINIMAC4 ^40^ can be used for pre-phasing and genotype imputation, respectively. In addition, GUIDANCE accepts one or multiple reference panels, allowing the integration of the results obtained from all panels by selecting for each variant the genotypes from the reference panel that provides the highest imputation accuracy according to the IMPUTE2 info score or MINIMAC2 r^2^ (Supplementary Figure 26). GUIDANCE also performs a post-imputation quality control to eliminate low-quality imputed variants under the basis of the IMPUTE2 info score or MINIMAC2 r^2^ and the MAF.

After genotype imputation and post-imputation quality control, GUIDANCE applies SNPTEST for association testing, where additive, dominant, recessive, heterodominant and genotype models can be analyzed. Here, the user can decide to include several covariates for the association test, such as principal components to adjust for population stratification, or any other confounders. GUIDANCE also allows testing for multiple phenotypes or for a single phenotype with different covariates in the same execution. After association testing, variants are filtered by the deviation from Hardy-Weinberg equilibrium (HWE) *p*-value. Finally, GUIDANCE generates summary reports for each trait with all the inheritance models tested in the association and the corresponding graphical representation, i.e., Manhattan and Quantile-Quantile (Q-Q) plots (Supplementary Figure 1-22), also providing a matrix identifying cross-phenotype associations (Supplementary Table 7).

GUIDANCE can be executed as a a standalone compact program or as independent modules (see Supplementary Figure 25 for a list of independent modules) to facilitate the use of GUIDANCE into existing frameworks.

Further details can be found in the configuration file from the GUIDANCE execution at http://cg.bsc.es/guidance. Specific documentation to use this framework is available at http://cg.bsc.es/guidance, as well as the source code and the pre-compiled binaries that are available in the “download” section.

### The Analysis of GERA cohort

#### GERA cohort Description

GERA cohort data was obtained through dbGaP under accession phs000674.v1.p1. For further information about the specific phenotypes (ICD-9-CM codes) included in GERA, please visit its website on dbGaP (https://www.ncbi.nlm.nih.gov/gap/). The Resource for Genetic Epidemiology Research on Aging (GERA) Cohort was created by a RC2 “Grand Opportunity” grant that was awarded to the Kaiser Permanente Research Program on Genes, Environment, and Health (RPGEH) and the UCSF Institute for Human Genetics (AG036607; Schaefer/Risch, PIs). The RC2 project enabled genome-wide SNP genotyping (GWAS) to be conducted on a cohort of over 100 K adults who were members of the Kaiser Permanente Medical Care Plan, Northern California Region (KPNC), and participating in its RPGEH. The resulting GERA cohort is composed of 42% of males, 58% of females, and ranges in age from 18 to over 100 years old with an average age of 63 years at the time of the RPGEH survey (2007). 19% of the individuals are from non-European ancestry, while 81% are described as white non-Hispanic participants. After an explicit requirement of consent by email, data from 78,486 participants was deposited in dbGaP, with similar demographic characteristics to those of the initial genotyped cohort.

#### Quality Control

A subset of 62,281 subjects of European ancestry underwent quality control analyses. A 3-step quality control protocol was applied using PLINK ^41, 42^, and included 2 stages of SNP removal and an intermediate stage of sample exclusion.

The exclusion criteria for genetic markers consisted of: proportion of missingness ≥ 0.05, HWE *p* ≤ 1 × 10^−20^ for all the cohort, and MAF < 0.001. This protocol for genetic markers was performed twice, before and after sample exclusion.

For the individuals, we considered the following exclusion criteria: gender discordance, subject relatedness (pairs with PI-HAT ≥ 0.125 from which we removed the individual with the highest proportion of missingness), sample call rates ≥ 0.02 and population structure showing more than 4 standard deviations within the distribution of the study population according to the first seven principal components (Supplementary Figure 27). After QC, 56,637 subjects remained for the analysis (Supplementary Table 1).

#### Analyzing GERA cohort using GUIDANCE

GUIDANCE pre-phased the genotypes to whole haplotypes with SHAPEIT2, and then performed genotype imputation with IMPUTE2 using 1000G phase 3, UK10K, GoNL, and HRC as reference panels. After filtering variants with an info score < 0.7 and a MAF < 0.001, we tested for association with additive, dominant, recessive, heterodominant and genotypic logistic regression using SNPTEST, and including seven derived principal components, sex and age as covariates. To maximize power and accuracy, we combined the association results from the four reference panels by choosing for each variant, the genotypes from the reference panel that provided the best IMPUTE2 info score. For chromosome X we restricted the analysis to non-pseudoautosomal (non-PAR) regions and stratified the association analysis by sex to account for hemizygosity for males, while for females, we followed an autosomal model. Finally, we excluded variants with HWE controls *p* < 1 × 10^−6^ in the final results.

#### Identification of known and new associated loci

After the association test, GUIDANCE provided a list of variants that passed the *p*-value threshold specified in the configuration file (i.e., p ≤ 5.0 × 10^−8^). Using the “IRanges” R package ^43^, all the genome-wide significant variants were collapsed into ranges (500 kb) that define each associated locus.

To distinguish between known or new associated regions, for each top variant we looked for any proxy variant with an LD *r*^2^ > 0.35 in the GWAS catalog (accession 5 September 2019) associated with the same phenotype or a related one (for example, bone mineral density, cholesterol levels or diastolic/systolic blood pressure phenotypes for osteoporosis, dyslipidemia or hypertension, respectively). HLA regions at chromosome 6 were excluded since the particularities of these regions required further detailed studies on their LD pattern. Proxies were selected using LDlink (https://ldlink.nci.nih.gov/) ^44^.

We defined an experiment-wide significant *p*-value cutoff of *p* < 2.0 × 10^−8^ by applying the Bonferroni correction for 2.5 effective test (5.0 × 10^−8^ / 2.5 effective test). This factor of 2.5 was obtained from a simulation study when four genetic models (additive, dominant, recessive and genotypic) are used ^45^ since the genetic models are not independent. However, a new simulation study including the heterodominant model should be done for a more accurate effective number of tests.

### Replication with UK Biobank

#### Phenotype Curation

UK Biobank participants agreed to provide detailed information about their lifestyle, environment and medical history, to donate biological samples (for genotyping and for biochemical assays), to undergo measures and to have their health followed (http://www.ukbiobank.ac.uk/).

When collecting and analyzing a wide range of phenotypes from the UK Biobank, a central challenge was the curation and harmonization of the vast array of categorizations, variable scalings, and follow-up responses. Fortunately, to this end, the PHEnome Scan ANalysis Tool (or PHESANT: https://github.com/MRCIEU/PHESANT) ^46^ performs much of the transformations and recodings required to generate meaningful, interpretable phenotypes.

We have made further adjustments based on user feedback, owing to the value of transparency in generating our phenotype guidelines. Applying these changes to the PHESANT source code, phenotypes were parsed using our modified version (github.com/astheeggeggs/PHESANT) on a virtual machine on the Google Cloud Platform.

We first restricted to the subset of European individuals, before passing the resultant phenotypic data to PHESANT. The ‘variable list’ file and ‘data-coding’ file, whose formats are defined in the original version of PHESANT were updated as new phenotypes were added in the latest UK Biobank release. Re-codings of variables, and inherent orderings of categorical variables, are defined in the ‘data-coding’ file. The ‘Excluded’ column of the ‘variable list’ file defines the collection of variables that we do not wish to interrogate.

A high level overview of the PHESANT pipeline, our defaults, and the associated short flags for the phenomescan.r code are displayed in Supplementary Figure 28. In addition to the inverse-rank normalization applied to the collection of continuous phenotypes, we also consider the raw version of the continuous phenotype, with no transformation applied to the data.

Curation of the ICD10 codes was carried out separately for computational efficiency. For the ICD10 phenotype, individuals are assigned a vector of ICD10 diagnoses. We truncated these codes to two digits, and assigned each individual to either case or control status for that ICD10 code in turn by checking if their vector contains that code. Throughout, we assumed the data contained no missingness, so the sum of cases and controls throughout was the number of individuals in our ‘European’ subset of the UK Biobank data. As in the PHESANT categorical (multiple) phenotypes, ICD10 code case/control phenotypes were removed if less than 50 individuals had the diagnosis.

#### Association testing and meta-analysis for UK Biobank phenotypes

We performed the association testing for the curated phenotypes as implemented in SNPTEST for additive, dominant, recessive, heterodominant and genotypic inheritance models, as it has been described in the “Analyzing GERA cohort using GUIDANCE” section. For all genotypic variants identified in the discovery stage, we assigned the recessive model after we identified it as the underlying model.

After the association testing, we filtered and ordered all the phenotypes based on the *p*-value for the best model of inheritance obtained from the GERA cohort analysis, with special consideration to equivalent phenotypes or related traits.

With the association testing results of both GERA cohort and UK Biobank, we meta-analyzed the results using METAL ^47^. We use the inverse variance-weighted fixed effect model for all the variants except for the rs557998486 variant associated with macular degeneration, since its *beta*, calculated with the “em” method from SNPTEST, was inflated. Therefore, we performed a sample size based meta-analysis, which converts the direction of the effect and the *p*-value into a z-score.

For biomarkers, only the results from the first visit were taken into account since less than 10% of the cases where present in the second visit.

#### Association testing and meta-analysis with FinnGen

We used SAIGE ^48^ for recessive association testing using sex, age, PC1-10 and batch as covariates. We analyzed FinnGen release 5 that contains 218,792 individuals with a median age 62.6 and a mean age 59.8.

For the cardiovascular disease endpoints, we meta-analyzed the results using “rmeta” R package ^49^. For macular degeneration, we meta-analyzed the results using METAL as described in the previous section.

#### Dominance deviation test

To detect genuine differences between additive and non-additive signals, we performed a dominance deviation test for all 93 autosomal genome-wide significant loci.

Dominance deviation was tested by a logistic regression analysis using PLINK (v1.90b6.9, www.cog-genomics.org/plink/1.9/). Sex, age and the first 7 PCs were included as covariates.

#### Definition of 99% credible set of *PELO* locus

For the *PELO* locus, the fraction of aggregated variants that have a 99% probability of containing the causal one was identified. The 99% credible set of variants for the region was defined with a Bayesian refinement approach ^50^, considering variants with an r^2^> 0.1 with the leading one.

For each variant within the *PELO* locus, the credible set provides a posterior probability of being the causal one ^50^. The approximate Bayes factor (*ABF*) for each variant was estimated as

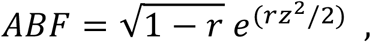

where

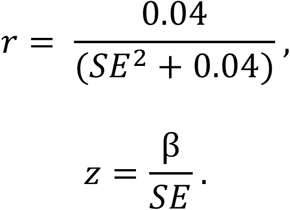

The β and the SE result from a logistic regression model testing for association. The posterior probability for each variant was calculated as

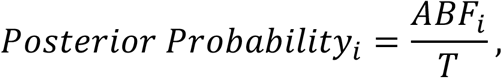

where *ABFi* corresponds to the approximate Bayes’ factor for the marker *i*, and *T* represents the sum of all the *ABF* values enclosed in the interval. As commonly employed by SNPTEST, this calculation assumes that the prior of the *β* is a Gaussian with mean 0 and variance 0.04.

Finally, the cumulative posterior probability was calculated after ranking the variants according to the *ABF* in decreasing order. Variants were included in the 99% credible set of the region until the cumulative posterior probability of association got over 0.99.

#### Gene expression and functional characterization

The eQTLGen Consortium (https://www.eqtlgen.org/cis-eqtls.html, last access on July 2019) and GTEx portal (https://gtexportal.org/, last access on July 2019) were used to find associations between our novel genetic associations and gene expression. When the variant was not available in the resources, a proxy SNP was used instead.

To determine whether any identified overlap between GERA GWAS loci and eQTLGen or GTEx eQTLs was due to a true shared association signal, we performed a colocalization analysis. Colocalization was assessed by a Bayesian test using summary statistics from both studies ^51^; summary statistics from the *cis* eQTLGen and GTEx were downloaded from the eQTLGen website and GTEx portal, respectively. The test was performed using the R package coloc v3.2-1 ^51, 52, 53^. The test provided a posterior probability for the GWAS locus and the eQTL to share the same causal variant(s).

We integrated available epigenomic datasets to examine the role of human pancreatic islet transcriptional regulation underlying rs77704739 association with type 2 diabetes. We used the WashU EpiGenome Browser (http://epigenomegateway.wustl.edu/browser/, last access on July 2019) and previously published RNA-seq, ATAC-seq and ChIP-seq assays of H3K4me3, H3K27ac, Mediator, CTCF and islet transcription factors (FOXA2, MAFB, NKX2.2, NKX6.1 and PDX1) in human pancreatic islets ^27, 28^ and islet regulome annotations ^28^.

## Supporting information

FinnGen Consortium banner

## Data Availability

The complete summary statistics are deposited at the Type 2 Diabetes Knowledge portal (www.type2diabetesgenetics.org/) and can be also accessed from http://cg.bsc.es/guidance. GUIDANCE is also available at http://cg.bsc.es/guidance.

## Acknowledgments

This work has been sponsored by the grant SEV-2011-00067 and SEV2015-0493 of Severo Ochoa Program, awarded by the Spanish Government, by the grant TIN2015-65316-P, awarded by the Spanish Ministry of Science and Innovation, and by the Generalitat de Catalunya (contract 2014-SGR-1051).This work was supported by an EFSD/Lilly research fellowship. Josep M. Mercader was supported by Sara Borrell Fellowship from the Instituto Carlos III. Sílvia Bonàs was FI-DGR Fellowship from FI-DGR 2013 from Agència de Gestió d’Ajuts Universitaris i de Recerca (AGAUR, Generalitat de Catalunya), and the American Diabetes Association Innovative and Clinical Translational Award 1-19-ICTS-068. Cecilia Salvoro received funding from the European Union’s Horizon 2020 research and innovation programme under the Marie Skłodowska-Curie grant agreement H2020-MSCA-COFUND-2016-754433. Cristian Ramon-Cortes pre-doctoral contract is financed by the Spanish Ministry of Science, Innovation, and Universities under contract BES-2016-076791. Elizabeth G. Atkinson was supported by the National Institutes of Mental Health (grants K01MH121659 and T32MH017119). Jose Florez was supported by NIH/NIDDK award K24 DK110550. This study made use of data generated by the UK10K Consortium, derived from samples from UK10K COHORT IMPUTATION (EGAS00001000713). A full list of the investigators who contributed to the generation of the data is available in www.UK10K.org. Funding for UK10K was provided by the Wellcome Trust under award WT091310. This study made use of data generated by the ‘Genome of the Netherlands’ project, which is funded by the Netherlands Organization for Scientific Research (grant no. 184021007). The data were made available as a Rainbow Project of BBMRI-NL. Samples were contributed by LifeLines (http://lifelines.nl/lifelines-research/general), the Leiden Longevity Study (http://www.healthy-ageing.nl; http://www.langleven.net), the Netherlands Twin Registry (NTR: http://www.tweelingenregister.org), the Rotterdam studies (http://www.erasmus-epidemiology.nl/rotterdamstudy) and the Genetic Research in Isolated Populations programme (http://www.epib.nl/research/geneticepi/research.html#gip). The sequencing was carried out in collaboration with the Beijing Institute for Genomics (BGI). This study also made use of data generated by The Haplotype Reference Consortium (HRC) accesed through The European Genome-phenome Archive at the European Bioinformatics Institute with the accession numbers EGAD00001002729, after a form agreed by the Barcelona Supercomputing Center (BSC) with WTSI. This research has been conducted using also the UK Biobank Resource (application number 31063 and 27892). The Genotype-Tissue Expression (GTEx) Project was supported by the Common Fund of the Office of the Director of the National Institutes of Health, and by NCI, NHGRI, NHLBI, NIDA, NIMH, and NINDS. The data used for the analyses described in this manuscript were obtained from the GTEx Portal on 07/16/2019. We acknowledge PRACE for awarding us access to both MareNostrum supercomputer from the Barcelona Supercomputing Center, based in Spain at Barcelona, and the SuperMUC supercomputer of the Leibniz Supercomputing Centre (LRZ), based in Garching at Germany (proposals numbers 2016143358 and 2016163985). The technical support group from the Barcelona Supercomputing Center is gratefully acknowledged. Finally, we thank all the Computational Genomics group at the BSC for their helpful discussions and valuable comments on the manuscript. We also acknowledge Elias Rodriguez Fos for designing the GUIDANCE logo.

## Authors Contributions

M.G-M., R.A., J.M.M., and D.T. conceived, planned, and performed the main analyses. M.G-M., J.M.M., and D.T. wrote the manuscript. M.G-M., R.A., M.P., C.R-C., F.S., J.E., C.D., E.T., and R.M.B. developed GUIDANCE. S.B-G. designed and performed the quality control. S.B-G and I. M-E. performed the functional characterization. C.S performed the dominance deviation test and the gene expression analysis. J.M.M., C.E.C., J.B.C, E.A., A.L., K.A., D.P., and J.C.F. contributed with UK Biobank data and analysis. S.R. and M.K. contributed with FinnGen data and analysis. J.M.M. and D.T. designed and supervised the study. All authors reviewed and approved the final manuscript.

