## Supplementary material for "The impact of non-additive genetic associations on age-related complex diseases": FinnGen Consortium banner

### Data Freeze 5:

### Contributors of FinnGen

##### Steering Committee


Aarno Palotie Institute for Molecular Medicine Finland, HiLIFE, University of Helsinki, Finland

Mark Daly Institute for Molecular Medicine Finland, HiLIFE, University of Helsinki, Finland


**Pharmaceutical companies**


Howard Jacob Abbvie, Chicago, IL, United States

Athena Matakidou Astra Zeneca, Cambridge, United Kingdom

Heiko Runz Biogen, Cambridge, MA, United States

Sally John Biogen, Cambridge, MA, United States

Robert Plenge Celgene, Summit, NJ, United States

Mark McCarthy Genentech, San Francisco, CA, United States

Julie Hunkapiller Genentech, San Francisco, CA, United States

Meg Ehm GlaxoSmithKline, Brentford, United Kingdom

Dawn Waterworth GlaxoSmithKline, Brentford, United Kingdom

Caroline Fox Merck, Kenilworth, NJ, United States

Anders Malarstig Pfizer, New York, NY, United States

Kathy Klinger Sanofi, Paris, France

Kathy Call Sanofi, Paris, France

Tim Behrens Maze Therapeutics, San Francisco, CA, United States

Patrick Loerch Janssen Biotech, Beerse, Belgium


**University of Helsinki & Biobanks**


Tomi Mäkelä HiLIFE, University of Helsinki, Finland, Finland

Jaakko Kaprio Institute for Molecular Medicine Finland, HiLIFE, Helsinki, Finland, Finland

Petri Virolainen Auria Biobank / University of Turku / Hospital District of Southwest Finland, Turku, Finland

Kari Pulkki Auria Biobank / University of Turku / Hospital District of Southwest Finland, Turku, Finland

Terhi Kilpi THL Biobank / The National Institute of Health and Welfare Helsinki, Finland

Markus Perola THL Biobank / The National Institute of Health and Welfare Helsinki, Finland

Jukka Partanen Finnish Red Cross Blood Service / Finnish Hematology Registry and Clinical Biobank, Helsinki, Finland

Anne Pitkäranta Helsinki Biobank / Helsinki University and Hospital District of Helsinki and Uusimaa, Helsinki

Riitta Kaarteenaho Northern Finland Biobank Borealis / University of Oulu / Northern Ostrobothnia Hospital District, Oulu, Finland

Seppo Vainio Northern Finland Biobank Borealis / University of Oulu / Northern Ostrobothnia Hospital District, Oulu, Finland

Miia Turpeinen Northern Finland Biobank Borealis / University of Oulu / Northern Ostrobothnia Hospital

Raisa Serpi Northern Finland Biobank Borealis / University of Oulu / Northern Ostrobothnia Hospital

Tarja Laitinen Finnish Clinical Biobank Tampere / University of Tampere / Pirkanmaa Hospital District, Tampere, Finland

Johanna Mäkelä Finnish Clinical Biobank Tampere / University of Tampere / Pirkanmaa Hospital District, Tampere, Finland

Veli-Matti Kosma Biobank of Eastern Finland / University of Eastern Finland / Northern Savo Hospital District, Kuopio, Finland

Urho Kujala Central Finland Biobank / University of Jyväskylä / Central Finland Health Care District, Jyväskylä, Finland

**Other Experts/ Non-Voting Members**

Outi Tuovila Business Finland, Helsinki, Finland

Minna Hendolin Business Finland, Helsinki, Finland

Raimo Pakkanen Business Finland, Helsinki, Finland

#### Scientific Committee

**Pharmaceutical companies**

Jeff Waring Abbvie, Chicago, IL, United States

Bridget Riley-Gillis Abbvie, Chicago, IL, United States

Athena Matakidou Astra Zeneca, Cambridge, United Kingdom

Heiko Runz Biogen, Cambridge, MA, United States

Jimmy Liu Biogen, Cambridge, MA, United States

Shameek Biswas Celgene, Summit, NJ, United States

Julie Hunkapiller Genentech, San Francisco, CA, United States

Dawn Waterworth GlaxoSmithKline, Brentford, United Kingdom

Meg Ehm GlaxoSmithKline, Brentford, United Kingdom

Dorothee Diogo Merck, Kenilworth, NJ, United States

Caroline Fox Merck, Kenilworth, NJ, United States

Anders Malarstig Pfizer, New York, NY, United States

Catherine Marshall Pfizer, New York, NY, United States

Xinli Hu Pfizer, New York, NY, United States

Kathy Call Sanofi, Paris, France

Kathy Klinger Sanofi, Paris, France

Matthias Gossel Sanofi, Paris, France

Robert Graham Maze Therapeutics, San Francisco, CA, United States

Tim Behrens Maze Therapeutics, San Francisco, CA, United States

Beryl Cummings Maze Therapeutics, San Francisco, CA, United States

Wilco Fleuren Janssen Biotech, Beerse, Belgium

**University of Helsinki & Biobanks**

Samuli Ripatti Institute for Molecular Medicine Finland, HiLIFE, Helsinki, Finland

Johanna Schleutker Auria Biobank / Univ. of Turku / Hospital District of Southwest Finland, Turku, Finland

Markus Perola THL Biobank / The National Institute of Health and Welfare Helsinki, Finland

Mikko Arvas Finnish Red Cross Blood Service / Finnish Hematology Registry and Clinical Biobank, Helsinki, Finland

Olli Carpen Helsinki Biobank / Helsinki University and Hospital District of Helsinki and Uusimaa, Helsinki

Reetta Hinttala Northern Finland Biobank Borealis / University of Oulu / Northern Ostrobothnia Hospital District, Oulu, Finland

Johannes Kettunen Northern Finland Biobank Borealis / University of Oulu / Northern Ostrobothnia Hospital District, Oulu, Finland

Johanna Mäkelä Finnish Clinical Biobank Tampere / University of Tampere / Pirkanmaa Hospital District, Tampere, Finland

Arto Mannermaa Biobank of Eastern Finland / University of Eastern Finland / Northern Savo Hospital District, Kuopio, Finland

Jari Laukkanen Central Finland Biobank / University of Jyväskylä / Central Finland Health Care District, Jyväskylä, Finland

Urho Kujala Central Finland Biobank / University of Jyväskylä / Central Finland Health Care District, Jyväskylä, Finland

**Other Experts/ Non-Voting Members**

| | |
|---|---|
| Outi Tuovila | Business Finland, Helsinki, Finland |
| Minna Hendolin | Business Finland, Helsinki, Finland |
| Raimo Pakkanen | Business Finland, Helsinki, Finland |

#### Clinical Groups

**Neurology Group**

| | |
|---|---|
| Hilkka Soininen | Northern Savo Hospital District, Kuopio, Finland |
| Valtteri Julkunen | Northern Savo Hospital District, Kuopio, Finland |
| Anne Remes | Northern Ostrobothnia Hospital District, Oulu, Finland |
| Reetta Kälviäinen | Northern Savo Hospital District, Kuopio, Finland |
| Mikko Hiltunen | Northern Savo Hospital District, Kuopio, Finland |
| Jukka Peltola | Pirkanmaa Hospital District, Tampere, Finland |
| Pentti Tienari | Hospital District of Helsinki and Uusimaa, Helsinki, Finland |
| Juha Rinne | Hospital District of Southwest Finland, Turku, Finland |
| Adam Ziemann | Abbvie, Chicago, IL, United States |
| Jeffrey Waring | Abbvie, Chicago, IL, United States |
| Sahar Esmaeeli | Abbvie, Chicago, IL, United States |
| Nizar Smaoui | Abbvie, Chicago, IL, United States |

| | |
|---|---|
| Anne Lehtonen | Abbvie, Chicago, IL, United States |
| Susan Eaton | Biogen, Cambridge, MA, United States |
| Heiko Runz | Biogen, Cambridge, MA, United States |
| Sanni Lahdenperä | Biogen, Cambridge, MA, United States |
| Janet van Adelsberg | Celgene, Summit, NJ, United States |
| Shameek Biswas | Celgene, Summit, NJ, United States |
| John Michon | Genentech, San Francisco, CA, United States |
| Geoff Kerchner | Genentech, San Francisco, CA, United States |
| Julie Hunkapiller | Genentech, San Francisco, CA, United States |
| Natalie Bowers | Genentech, San Francisco, CA, United States |
| Edmond Teng | Genentech, San Francisco, CA, United States |
| John Eicher | Merck, Kenilworth, NJ, United States |
| Vinay Mehta | Merck, Kenilworth, NJ, United States |
| Padhraig Gormley | Merck, Kenilworth, NJ, United States |
| Kari Linden | Pfizer, New York, NY, United States |
| Christopher Whelan | Pfizer, New York, NY, United States |
| Fanli Xu | GlaxoSmithKline, Brentford, United Kingdom |
| David Pulford | GlaxoSmithKline, Brentford, United Kingdom |

**Gastroenterology Group**

| | |
|---|---|
| Martti Färkkilä | Hospital District of Helsinki and Uusimaa, Helsinki, Finland |
| Sampsa Pikkarainen | Hospital District of Helsinki and Uusimaa, Helsinki, Finland |
| Airi Jussila | Pirkanmaa Hospital District, Tampere, Finland |
| Timo Blomster | Northern Ostrobothnia Hospital District, Oulu, Finland |
| Mikko Kiviniemi | Northern Savo Hospital District, Kuopio, Finland |
| Markku Voutilainen | Hospital District of Southwest Finland, Turku, Finland |
| Bob Georgantas | Abbvie, Chicago, IL, United States |
| Graham Heap | Abbvie, Chicago, IL, United States |
| Jeffrey Waring | Abbvie, Chicago, IL, United States |
| Nizar Smaoui | Abbvie, Chicago, IL, United States |

| | |
|---|---|
| Fedik Rahimov | Abbvie, Chicago, IL, United States |
| Anne Lehtonen | Abbvie, Chicago, IL, United States |
| Keith Usiskin | Celgene, Summit, NJ, United States |
| Tim Lu | Genentech, San Francisco, CA, United States |
| Natalie Bowers | Genentech, San Francisco, CA, United States |
| Danny Oh | Genentech, San Francisco, CA, United States |
| John Michon | Genentech, San Francisco, CA, United States |
| Vinay Mehta | Merck, Kenilworth, NJ, United States |
| Kirsi Kalpala | Pfizer, New York, NY, United States |
| Melissa Miller | Pfizer, New York, NY, United States |
| Xinli Hu | Pfizer, New York, NY, United States |
| Linda McCarthy | GlaxoSmithKline, Brentford, United Kingdom |

**Rheumatology Group**

| | |
|---|---|
| Kari Eklund | Hospital District of Helsinki and Uusimaa, Helsinki, Finland |
| Antti Palomäki | Hospital District of Southwest Finland, Turku, Finland |
| Pia Isomäki | Pirkanmaa Hospital District, Tampere, Finland |
| Laura Pirilä | Hospital District of Southwest Finland, Turku, Finland |
| Oili Kaipiainen-Seppänen | Northern Savo Hospital District, Kuopio, Finland |
| Johanna Huhtakangas | Northern Ostrobothnia Hospital District, Oulu, Finland |
| Bob Georgantas | Abbvie, Chicago, IL, United States |
| Jeffrey Waring | Abbvie, Chicago, IL, United States |
| Fedik Rahimov | Abbvie, Chicago, IL, United States |
| Apinya Lertratanakul | Abbvie, Chicago, IL, United States |
| Nizar Smaoui | Abbvie, Chicago, IL, United States |
| Anne Lehtonen | Abbvie, Chicago, IL, United States |
| David Close | Astra Zeneca, Cambridge, United Kingdom |
| Marla Hochfeld | Celgene, Summit, NJ, United States |
| Natalie Bowers | Genentech, San Francisco, CA, United States |
| John Michon | Genentech, San Francisco, CA, United States |

| | |
|---|---|
| Dorothee Diogo | Merck, Kenilworth, NJ, United States |
| Vinay Mehta | Merck, Kenilworth, NJ, United States |
| Kirsi Kalpala | Pfizer, New York, NY, United States |
| Nan Bing | Pfizer, New York, NY, United States |
| Xinli Hu | Pfizer, New York, NY, United States |
| Jorge Esparza Gordillo | GlaxoSmithKline, Brentford, United Kingdom |
| Nina Mars | Institute for Molecular Medicine Finland, HiLIFE, Helsinki, Finland |

**Pulmonology Group**

| | |
|---|---|
| Tarja Laitinen | Pirkanmaa Hospital District, Tampere, Finland |
| Margit Pelkonen | Northern Savo Hospital District, Kuopio, Finland |
| Paula Kauppi | Hospital District of Helsinki and Uusimaa, Helsinki, Finland |
| Hannu Kankaanranta | Pirkanmaa Hospital District, Tampere, Finland |
| Terttu Harju | Northern Ostrobothnia Hospital District, Oulu, Finland |
| Nizar Smaoui | Abbvie, Chicago, IL, United States |
| David Close | Astra Zeneca, Cambridge, United Kingdom |
| Susan Eaton | Biogen, Cambridge, MA, United States |
| Steven Greenberg | Celgene, Summit, NJ, United States |
| Hubert Chen | Genentech, San Francisco, CA, United States |
| Natalie Bowers | Genentech, San Francisco, CA, United States |
| John Michon | Genentech, San Francisco, CA, United States |
| Vinay Mehta | Merck, Kenilworth, NJ, United States |
| Jo Betts | GlaxoSmithKline, Brentford, United Kingdom |
| Soumitra Ghosh | GlaxoSmithKline, Brentford, United Kingdom |

**Cardiometabolic Diseases Group**

| | |
|---|---|
| Veikko Salomaa | The National Institute of Health and Welfare Helsinki, Finland |
| Teemu Niiranen | The National Institute of Health and Welfare Helsinki, Finland |
| Markus Juonala | Hospital District of Southwest Finland, Turku, Finland |
| Kaj Metsärinne | Hospital District of Southwest Finland, Turku, Finland |

| | |
|---|---|
| Mika Kähönen | Pirkanmaa Hospital District, Tampere, Finland |
| Juhani Junttila | Northern Ostrobothnia Hospital District, Oulu, Finland |
| Markku Laakso | Northern Savo Hospital District, Kuopio, Finland |
| Jussi Pihlajamäki | Northern Savo Hospital District, Kuopio, Finland |
| Juha Sinisalo | Hospital District of Helsinki and Uusimaa, Helsinki, Finland |
| Marja-Riitta Taskinen | Hospital District of Helsinki and Uusimaa, Helsinki, Finland |
| Tiinamaija Tuomi | Hospital District of Helsinki and Uusimaa, Helsinki, Finland |
| Jari Laukkanen | Central Finland Health Care District, Jyväskylä, Finland |
| Ben Challis | Astra Zeneca, Cambridge, United Kingdom |
| Andrew Peterson | Genentech, San Francisco, CA, United States |
| Julie Hunkapiller | Genentech, San Francisco, CA, United States |
| Natalie Bowers | Genentech, San Francisco, CA, United States |
| John Michon | Genentech, San Francisco, CA, United States |
| Dorothee Diogo | Merck, Kenilworth, NJ, United States |
| Audrey Chu | Merck, Kenilworth, NJ, United States |
| Vinay Mehta | Merck, Kenilworth, NJ, United States |
| Jaakko Parkkinen | Pfizer, New York, NY, United States |
| Melissa Miller | Pfizer, New York, NY, United States |
| Anthony Muslin | Sanofi, Paris, France |
| Dawn Waterworth | GlaxoSmithKline, Brentford, United Kingdom |

**Oncology Group**

| | |
|---|---|
| Heikki Joensuu | Hospital District of Helsinki and Uusimaa, Helsinki, Finland |
| Olli Carpen | Hospital District of Helsinki and Uusimaa, Helsinki, Finland |
| Tuomo Meretoja | Hospital District of Helsinki and Uusimaa, Helsinki, Finland |
| Lauri Aaltonen | Hospital District of Helsinki and Uusimaa, Helsinki, Finland |
| Johanna Mattson | Hospital District of Helsinki and Uusimaa, Helsinki, Finland |
| Johanna Schleutker | University of Turku, Turku, Finland |

| | |
|---|---|
| Annika Auranen | Pirkanmaa Hospital District , Tampere, Finland |

| | |
|---|---|
| Peeter Karihtala | Northern Ostrobothnia Hospital District, Oulu, Finland |
| Saila Kauppila | Northern Ostrobothnia Hospital District, Oulu, Finland |
| Päivi Auvinen | Northern Savo Hospital District, Kuopio, Finland |
| Klaus Elenius | Hospital District of Southwest Finland, Turku, Finland |
| Relja Popovic | Abbvie, Chicago, IL, United States |
| Jeffrey Waring | Abbvie, Chicago, IL, United States |
| Bridget Riley-Gillis | Abbvie, Chicago, IL, United States |
| Anne Lehtonen | Abbvie, Chicago, IL, United States |
| Athena Matakidou | Astra Zeneca, Cambridge, United Kingdom |
| Jennifer Schutzman | Genentech, San Francisco, CA, United States |
| Julie Hunkapiller | Genentech, San Francisco, CA, United States |
| Natalie Bowers | Genentech, San Francisco, CA, United States |
| John Michon | Genentech, San Francisco, CA, United States |
| Vinay Mehta | Merck, Kenilworth, NJ, United States |
| Andrey Loboda | Merck, Kenilworth, NJ, United States |
| Aparna Chhibber | Merck, Kenilworth, NJ, United States |
| Heli Lehtonen | Pfizer, New York, NY, United States |
| Stefan McDonough | Pfizer, New York, NY, United States |
| Marika Crohns | Sanofi, Paris, France |
| Diptee Kulkarni | GlaxoSmithKline, Brentford, United Kingdom¨ |

**Opthalmology Group**

| | |
|---|---|
| Kai Kaarniranta | Northern Savo Hospital District, Kuopio, Finland |
| Joni A Turunen | Hospital District of Helsinki and Uusimaa, Helsinki, Finland |
| Terhi Ollila | Hospital District of Helsinki and Uusimaa, Helsinki, Finland |
| Sanna Seitsonen | Hospital District of Helsinki and Uusimaa, Helsinki, Finland |
| Hannu Uusitalo | Pirkanmaa Hospital District, Tampere, Finland |
| Vesa Aaltonen | Hospital District of Southwest Finland, Turku, Finland |
| Hannele Uusitalo-Järvinen | Pirkanmaa Hospital District, Tampere, Finland |
| Marja Luodonpää | Northern Ostrobothnia Hospital District, Oulu, Finland |

| | |
|---|---|
| Nina Hautala | Northern Ostrobothnia Hospital District, Oulu, Finland |
| Heiko Runz | Biogen, Cambridge, MA, United States |
| Stephanie Loomis | Biogen, Cambridge, MA, United States |
| Erich Strauss | Genentech, San Francisco, CA, United States |
| Natalie Bowers | Genentech, San Francisco, CA, United States |
| Hao Chen | Genentech, San Francisco, CA, United States |
| John Michon | Genentech, San Francisco, CA, United States |
| Anna Podgornaia | Merck, Kenilworth, NJ, United States |
| Vinay Mehta | Merck, Kenilworth, NJ, United States |
| Dorothee Diogo | Merck, Kenilworth, NJ, United States |
| Joshua Hoffman | GlaxoSmithKline, Brentford, United Kingdom |

**Dermatology Group**

| | |
|---|---|
| Kaisa Tasanen | Northern Ostrobothnia Hospital District, Oulu, Finland |
| Laura Huilaja | Northern Ostrobothnia Hospital District, Oulu, Finland |
| Katariina Hannula-Jouppi | Hospital District of Helsinki and Uusimaa, Helsinki, Finland |
| Teea Salmi | Pirkanmaa Hospital District, Tampere, Finland |
| Sirkku Peltonen | Hospital District of Southwest Finland, Turku, Finland |
| Leena Koulu | Hospital District of Southwest Finland, Turku, Finland |
| Ilkka Harvima | Northern Savo Hospital District, Kuopio, Finland |
| Kirsi Kalpala | Pfizer, New York, NY, United States |
| Ying Wu | Pfizer, New York, NY, United States |
| David Choy | Genentech, San Francisco, CA, United States |
| John Michon | Genentech, San Francisco, CA, United States |
| Nizar Smaoui | Abbvie, Chicago, IL, United States |
| Fedik Rahimov | Abbvie, Chicago, IL, United States |
| Anne Lehtonen | Abbvie, Chicago, IL, United States |
| Dawn Waterworth | GlaxoSmithKline, Brentford, United Kingdom |

**Odontology Group**

Pirkko Pussinen Hospital District of Helsinki and Uusimaa, Helsinki, Finland

Aino Salminen Hospital District of Helsinki and Uusimaa, Helsinki, Finland

Tuula Salo Hospital District of Helsinki and Uusimaa, Helsinki, Finland

David Rice Hospital District of Helsinki and Uusimaa, Helsinki, Finland

Pekka Nieminen Hospital District of Helsinki and Uusimaa, Helsinki, Finland

Ulla Palotie Hospital District of Helsinki and Uusimaa, Helsinki, Finland

Maria Siponen Northern Savo Hospital District, Kuopio, Finland

Liisa Suominen Northern Savo Hospital District, Kuopio, Finland

Päivi Mäntylä Northern Savo Hospital District, Kuopio, Finland

Ulvi Gursoy Hospital District of Southwest Finland, Turku, Finland

Vuokko Anttonen Northern Ostrobothnia Hospital District, Oulu, Finland

Kirsi Sipilä Northern Ostrobothnia Hospital District, Oulu, Finland

#### FinnGen Analysis working group

##### Analysis Group

Justin Wade Davis Abbvie, Chicago, IL, United States

Bridget Riley-Gillis Abbvie, Chicago, IL, United States

Danjuma Quarless Abbvie, Chicago, IL, United States

Fedik Rahimov Abbvie, Chicago, IL, United States

Sahar Esmaeeli Abbvie, Chicago, IL, United States

Slavé Petrovski Astra Zeneca, Cambridge, United Kingdom

Eleonor Wigmore Astra Zeneca, Cambridge, United Kingdom

Jimmy Liu Biogen, Cambridge, MA, United States

Chia-Yen Chen Biogen, Cambridge, MA, United States

Paola Bronson Biogen, Cambridge, MA, United States

Ellen Tsai Biogen, Cambridge, MA, United States

Stephanie Loomis Biogen, Cambridge, MA, United States

Yunfeng Huang Biogen, Cambridge, MA, United States

Joseph Maranville Celgene, Summit, NJ, United States

Shameek Biswas Celgene, Summit, NJ, United States

Elmutaz Shaikho Elhaj Mohammed Celgene, Summit, NJ, United States

Samir Wadhawan Bristol-Meyers-Squibb

| | |
|---|---|
| Erika Kvikstad | Bristol-Meyers-Squibb |
| Minal Caliskan | Bristol-Meyers-Squibb |
| Diana Chang | Genentech, San Francisco, CA, United States |
| Julie Hunkapiller | Genentech, San Francisco, CA, United States |
| Tushar Bhangale | Genentech, San Francisco, CA, United States |
| Natalie Bowers | Genentech, San Francisco, CA, United States |
| Sarah Pendergrass | Genentech, San Francisco, CA, United States |
| Dorothee Diogo | Merck, Kenilworth, NJ, United States |
| Emily Holzinger | Merck, Kenilworth, NJ, United States |
| Padhraig Gormley | Merck, Kenilworth, NJ, United States |
| Xing Chen | Pfizer, New York, NY, United States |
| Åsa Hedman | Pfizer, New York, NY, United States |
| Karen S King | GlaxoSmithKline, Brentford, United Kingdom |
| Clarence Wang | Sanofi, Paris, France |
| Ethan Xu | Sanofi, Paris, France |
| Franck Auge | Sanofi, Paris, France |
| Clement Chatelain | Sanofi, Paris, France |
| Deepak Rajpal | Sanofi, Paris, France |
| Dongyu Liu | Sanofi, Paris, France |
| Katherine Call | Sanofi, Paris, France |
| Tai-he Xia | Sanofi, Paris, France |
| Beryl Cummings | Maze Therapeutics, San Francisco, CA, United States |
| Matt Brauer | Maze Therapeutics, San Francisco, CA, United States |
| Mitja Kurki | Institute for Molecular Medicine Finland, HiLIFE, University of Helsinki, Finland / Broad Institute, Cambridge, MA, United States |
| Samuli Ripatti | Institute for Molecular Medicine Finland, HiLIFE, University of Helsinki, Finland |
| Mark Daly | Institute for Molecular Medicine Finland, HiLIFE, University of Helsinki, Finland |
| Juha Karjalainen | Institute for Molecular Medicine Finland, HiLIFE, University of Helsinki, Finland / Broad Institute, Cambridge, MA, United States |
| Aki Havulinna | Institute for Molecular Medicine Finland, HiLIFE, University of Helsinki, Finland |

Anu Jalanko Institute for Molecular Medicine Finland, HiLIFE, University of Helsinki, Finland

Priit Palta Institute for Molecular Medicine Finland, HiLIFE, University of Helsinki, Finland

Pietro della Briotta Parolo Institute for Molecular Medicine Finland, HiLIFE, University of Helsinki, Finland

Wei Zhou Broad Institute, Cambridge, MA, United States

Susanna Lemmelä Institute for Molecular Medicine Finland, HiLIFE, University of Helsinki, Finland

Manuel Rivas University of Stanford, Stanford, CA, United States

Jarmo Harju Institute for Molecular Medicine Finland, HiLIFE, University of Helsinki, Finland

Aarno Palotie Institute for Molecular Medicine Finland, HiLIFE, University of Helsinki, Finland

Arto Lehisto Institute for Molecular Medicine Finland, HiLIFE, University of Helsinki, Finland

Andrea Ganna Institute for Molecular Medicine Finland, HiLIFE, University of Helsinki, Finland

Vincent Llorens Institute for Molecular Medicine Finland, HiLIFE, University of Helsinki, Finland

Hannele Laivuori Institute for Molecular Medicine Finland, HiLIFE, University of Helsinki, Finland

Sina Rüeger Institute for Molecular Medicine Finland, HiLIFE, University of Helsinki, Finland

Mari E Niemi Institute for Molecular Medicine Finland, HiLIFE, University of Helsinki, Finland

Taru Tukiainen Institute for Molecular Medicine Finland, HiLIFE, University of Helsinki, Finland

Mary Pat Reeve Institute for Molecular Medicine Finland, HiLIFE, University of Helsinki, Finland

Henrike Heyne Institute for Molecular Medicine Finland, HiLIFE, University of Helsinki, Finland

Nina Mars Institute for Molecular Medicine Finland, HiLIFE, University of Helsinki, Finland

Kimmo Palin University of Helsinki, Helsinki, Finland

Javier Garcia-Tabuenca University of Tampere, Tampere, Finland

Harri Siirtola University of Tampere, Tampere, Finland

Tuomo Kiiskinen Institute for Molecular Medicine Finland, HiLIFE, University of Helsinki, Finland

Jiwoo Lee Institute for Molecular Medicine Finland, HiLIFE, University of Helsinki, Finland / Broad Institute, Cambridge, MA, United States

Kristin Tsuo Institute for Molecular Medicine Finland, HiLIFE, University of Helsinki, Finland / Broad Institute, Cambridge, MA, United States

Amanda Elliott Institute for Molecular Medicine Finland, HiLIFE, University of Helsinki, Finland / Broad Institute, Cambridge, MA, United States

Kati Kristiansson THL Biobank / The National Institute of Health and Welfare Helsinki, Finland

Mikko Arvas Finnish Red Cross Blood Service, Helsinki, Finland

Kati Hyvärinen Finnish Red Cross Blood Service, Helsinki, Finland

Jarmo Ritari Finnish Red Cross Blood Service, Helsinki, Finland

Miika Koskinen Helsinki Biobank / Helsinki University and Hospital District of Helsinki and Uusimaa, Helsinki

Olli Carpen Helsinki Biobank / Helsinki University and Hospital District of Helsinki and Uusimaa, Helsinki

Johannes Kettunen Northern Finland Biobank Borealis / University of Oulu / Northern Ostrobothnia Hospital District, Oulu, Finland

Katri Pylkäs Northern Finland Biobank Borealis / University of Oulu / Northern Ostrobothnia Hospital District, Oulu, Finland

Marita Kalaoja Northern Finland Biobank Borealis / University of Oulu / Northern Ostrobothnia Hospital District, Oulu, Finland

Minna Karjalainen Northern Finland Biobank Borealis / University of Oulu / Northern Ostrobothnia Hospital District, Oulu, Finland

Tuomo Mantere Northern Finland Biobank Borealis / University of Oulu / Northern Ostrobothnia Hospital District, Oulu, Finland

Eeva Kangasniemi Finnish Clinical Biobank Tampere / University of Tampere / Pirkanmaa Hospital District, Tampere, Finland

Sami Heikkinen Biobank of Eastern Finland / University of Eastern Finland / Northern Savo Hospital District, Kuopio, Finland

Arto Mannermaa Biobank of Eastern Finland / University of Eastern Finland / Northern Savo Hospital District, Kuopio, Finland

Eija Laakkonen Central Finland Biobank / University of Jyväskylä / Central Finland Health Care District, Jyväskylä, Finland

Juha Kononen Central Finland Biobank / University of Jyväskylä / Central Finland Health Care District, Jyväskylä, Finland

Csilla Sipeky University of Turku, Turku, Finland

Samuel Heron University of Turku, Turku, Finland

Antti Karlsson Auria Biobank / University of Turku / Hospital District of Southwest Finland, Turku, Finland

Dhanaprakash Jambulingam University of Turku, Turku, Finland

Venkat Subramaniam Rathinakannan University of Turku, Turku, Finland

#### Biobank directors

Lila Kallio Auria Biobank / University of Turku / Hospital District of Southwest Finland, Turku, Finland

Sirpa Soini THL Biobank / The National Institute of Health and Welfare Helsinki, Finland

Jukka Partanen Finnish Red Cross Blood Service / Finnish Hematology Registry and Clinical Biobank, Helsinki, Finland

Eero Punkka Helsinki Biobank / Helsinki University and Hospital District of Helsinki and Uusimaa, Helsinki

Raisa Serpi Northern Finland Biobank Borealis / University of Oulu / Northern Ostrobothnia Hospital District, Oulu, Finland

Johanna Mäkelä Finnish Clinical Biobank Tampere / University of Tampere / Pirkanmaa Hospital District, Tampere, Finland

Veli-Matti Kosma Biobank of Eastern Finland / University of Eastern Finland / Northern Savo Hospital District, Kuopio, Finland

Teijo Kuopio Central Finland Biobank / University of Jyväskylä / Central Finland Health Care District, Jyväskylä, Finland

#### FinnGen Teams

##### Administration

Anu Jalanko Institute for Molecular Medicine Finland, HiLIFE, University of Helsinki, Finland

Risto Kajanne Institute for Molecular Medicine Finland, HiLIFE, University of Helsinki, Finland

Mervi Aavikko Institute for Molecular Medicine Finland, HiLIFE, University of Helsinki, Finland

Manuel González Jiménez Institute for Molecular Medicine Finland, HiLIFE, University of Helsinki, Finland

##### Analysis

Mitja Kurki Institute for Molecular Medicine Finland, HiLIFE, University of Helsinki, Finland / Broad Institute, Cambridge, MA, United States

Juha Karjalainen Institute for Molecular Medicine Finland, HiLIFE, University of Helsinki, Finland / Broad Institute, Cambridge, MA, United States

Pietro della Briotta Parola Institute for Molecular Medicine Finland, HiLIFE, University of Helsinki, Finland

Sina Rüeger Institute for Molecular Medicine Finland, HiLIFE, University of Helsinki, Finland

Arto Lehistö Institute for Molecular Medicine Finland, HiLIFE, University of Helsinki, Finland

Wei Zhou Broad Institute, Cambridge, MA, United States

Masahiro Kanai Broad Institute, Cambridge, MA, United States

**Clinical Endpoint Development**

Hannele Laivuori Institute for Molecular Medicine Finland, HiLIFE, University of Helsinki, Finland

Aki Havulinna Institute for Molecular Medicine Finland, HiLIFE, University of Helsinki, Finland

Susanna Lemmelä Institute for Molecular Medicine Finland, HiLIFE, University of Helsinki, Finland

Tuomo Kiiskinen Institute for Molecular Medicine Finland, HiLIFE, University of Helsinki, Finland

**Communication**

Mari Kaunisto Institute for Molecular Medicine Finland, HiLIFE, University of Helsinki, Finland

**Data Management and IT Infrastructure**

Jarmo Harju Institute for Molecular Medicine Finland, HiLIFE, University of Helsinki, Finland

Elina Kilpeläinen Institute for Molecular Medicine Finland, HiLIFE, University of Helsinki, Finland

Timo P. Sipilä Institute for Molecular Medicine Finland, HiLIFE, University of Helsinki, Finland

Georg Brein Institute for Molecular Medicine Finland, HiLIFE, University of Helsinki, Finland

Oluwaseun A. Dada Institute for Molecular Medicine Finland, HiLIFE, University of Helsinki, Finland

Ghazal Awaisa Institute for Molecular Medicine Finland, HiLIFE, University of Helsinki, Finland

Anastasia Shcherban Institute for Molecular Medicine Finland, HiLIFE, University of Helsinki, Finland

**Genotyping**

Kati Donner Institute for Molecular Medicine Finland, HiLIFE, University of Helsinki, Finland

Timo P. Sipilä Institute for Molecular Medicine Finland, HiLIFE, University of Helsinki, Finland

**Sample Collection Coordination**

Anu Loukola Helsinki Biobank / Helsinki University and Hospital District of Helsinki and Uusimaa, Helsinki

**Sample Logistics**

Päivi Laiho THL Biobank / The National Institute of Health and Welfare Helsinki, Finland

Tuuli Sistonen THL Biobank / The National Institute of Health and Welfare Helsinki, Finland

Essi Kaiharju THL Biobank / The National Institute of Health and Welfare Helsinki, Finland

Markku Laukkanen THL Biobank / The National Institute of Health and Welfare Helsinki, Finland

Elina Järvensivu THL Biobank / The National Institute of Health and Welfare Helsinki, Finland

Sini Lähteenmäki THL Biobank / The National Institute of Health and Welfare Helsinki, Finland

Lotta Männikkö THL Biobank / The National Institute of Health and Welfare Helsinki, Finland

Regis Wong THL Biobank / The National Institute of Health and Welfare Helsinki, Finland

**Registry Data Operations**

Hannele Mattsson THL Biobank / The National Institute of Health and Welfare Helsinki, Finland

Kati Kristiansson THL Biobank / The National Institute of Health and Welfare Helsinki, Finland

Susanna Lemmelä Institute for Molecular Medicine Finland, HiLIFE, University of Helsinki, Finland

Tero Hiekkalinna THL Biobank / The National Institute of Health and Welfare Helsinki, Finland

Teemu Paajanen THL Biobank / The National Institute of Health and Welfare Helsinki, Finland

**Sequencing Informatics**

Priit Palta Institute for Molecular Medicine Finland, HiLIFE, University of Helsinki, Finland

Kalle Pärn Institute for Molecular Medicine Finland, HiLIFE, University of Helsinki, Finland

**Trajectory Team**

Tarja Laitinen Pirkanmaa Hospital District, Tampere, Finland

Harri Siirtola University of Tampere, Tampere, Finland

Javier Gracia-Tabuenca University of Tampere, Tampere, Finland
