## Supplementary Figure 1-22 for "The impact of non-additive genetic associations on age-related complex diseases"

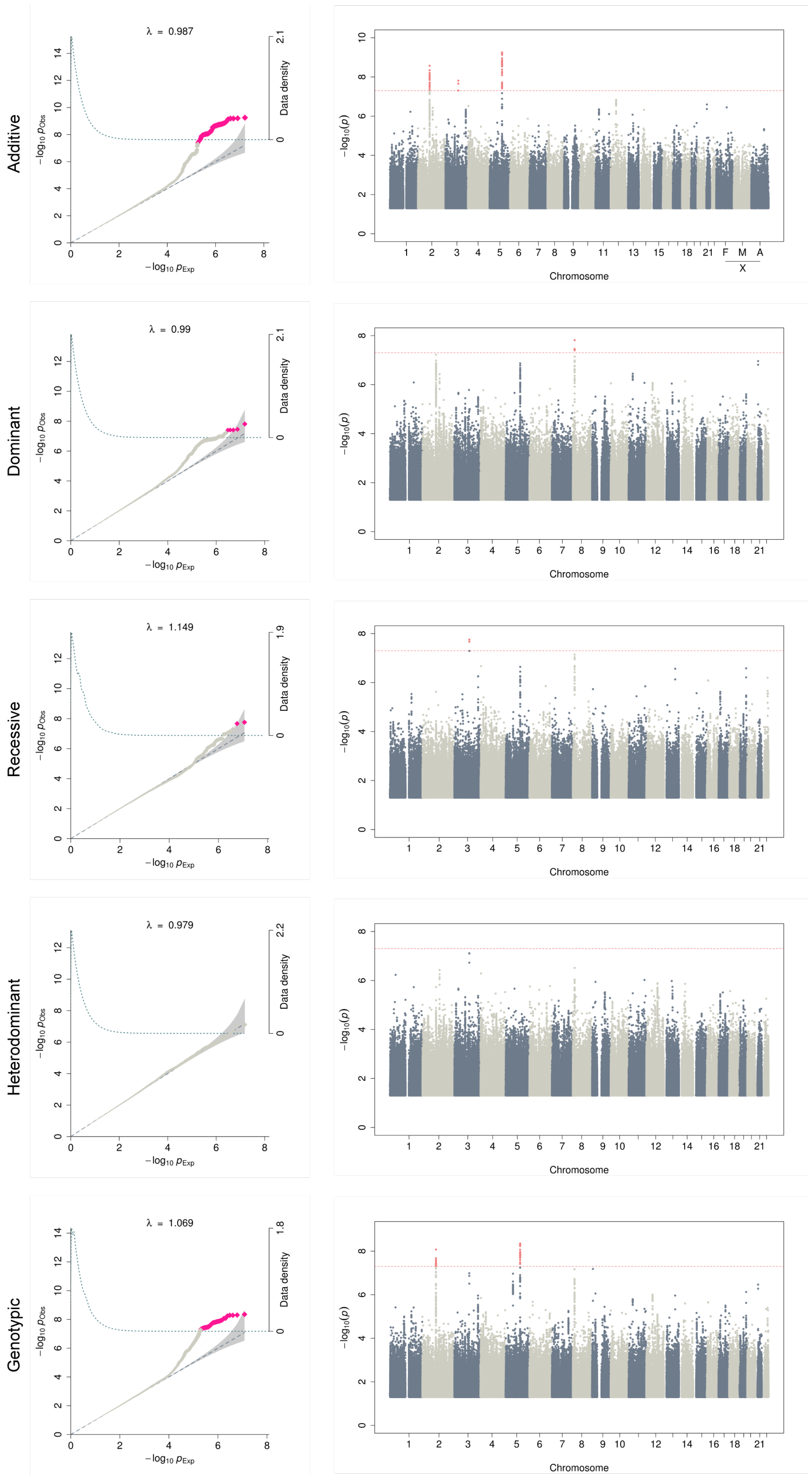


**Supplementary Figure 1. Q-Q plots and Manhattan plots for allergic rhinitis.**

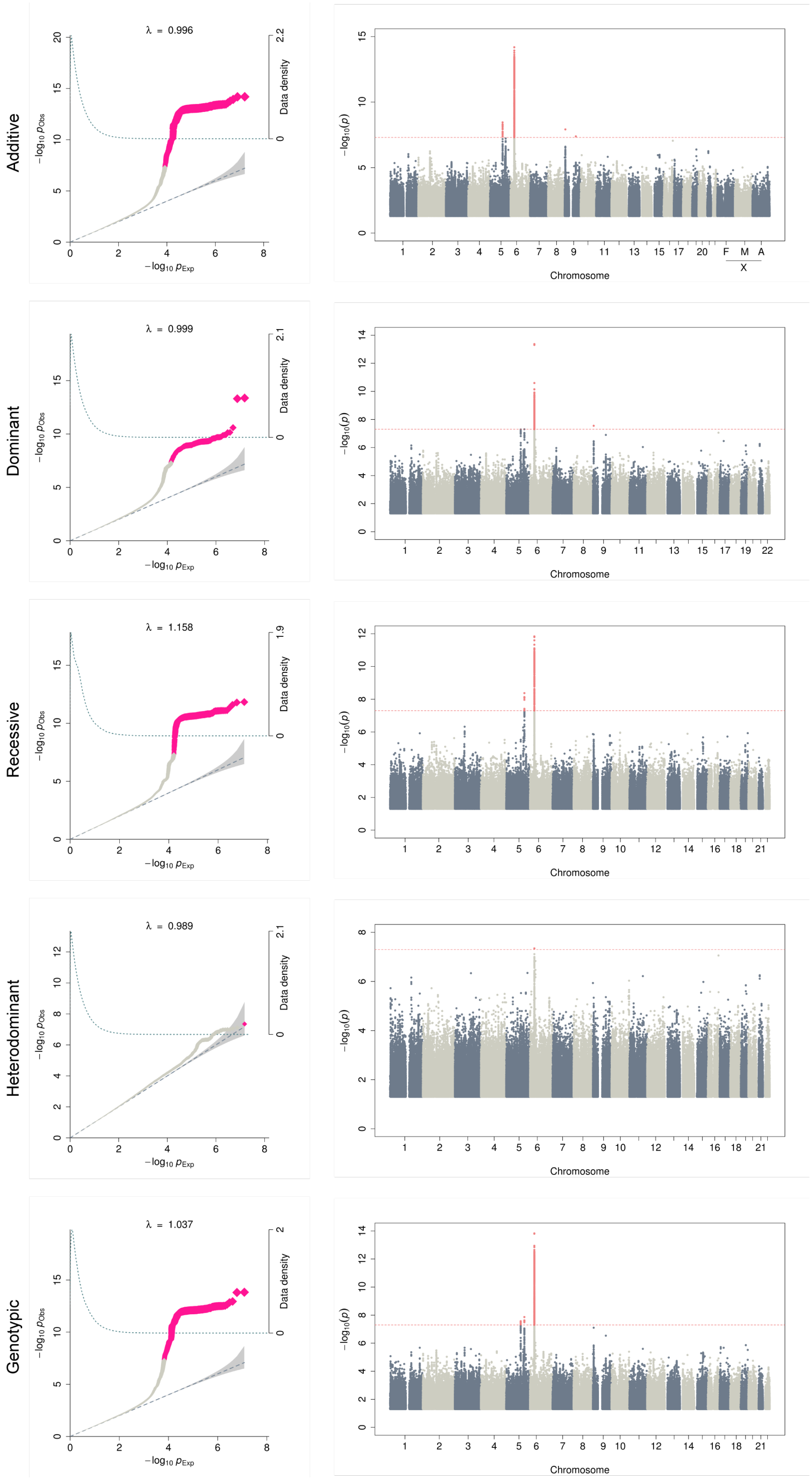


**Supplementary Figure 2. Q-Q plots and Manhattan plots for asthma.**

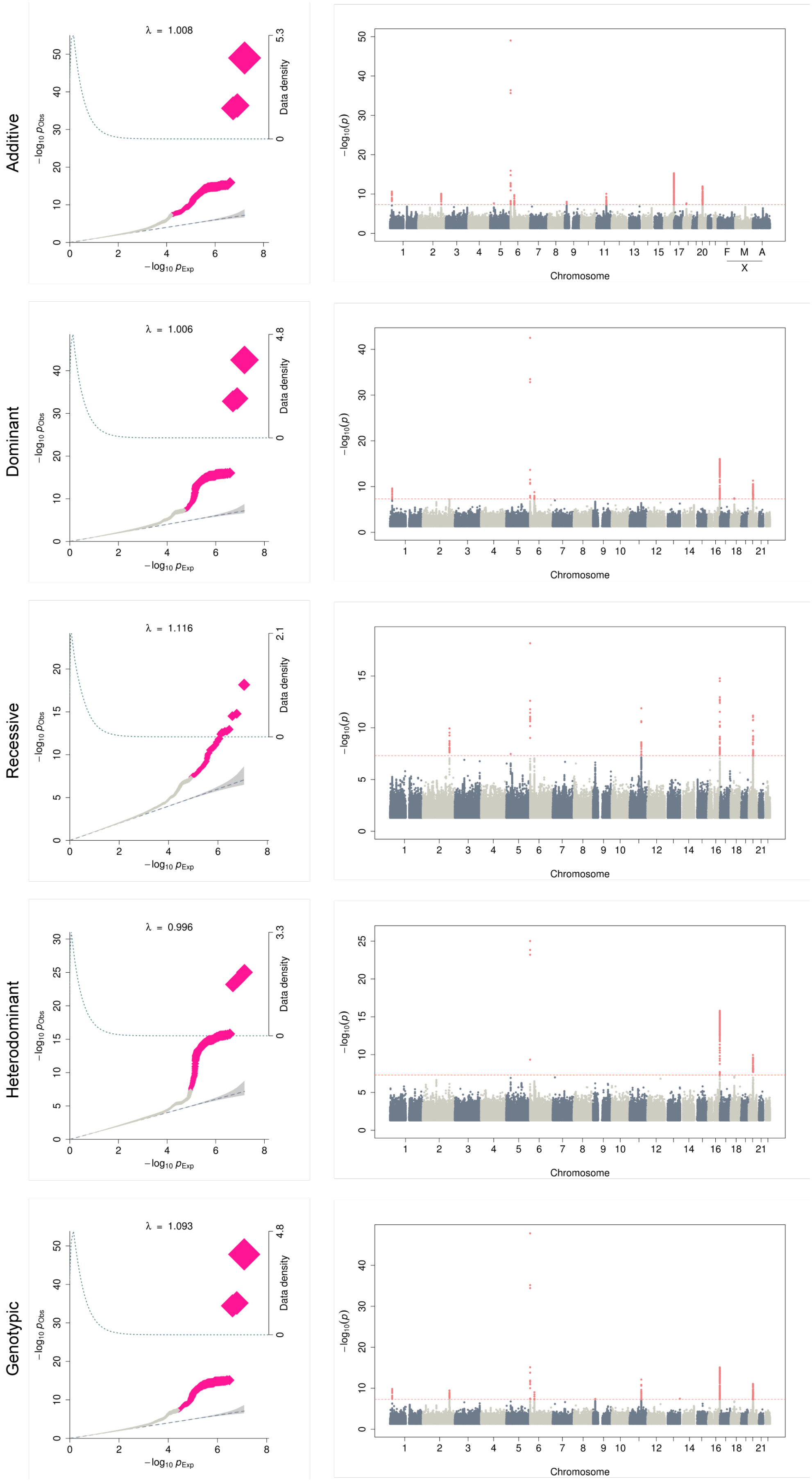


**Supplementary Figure 3. Q-Q plots and Manhattan plots for cancer.**

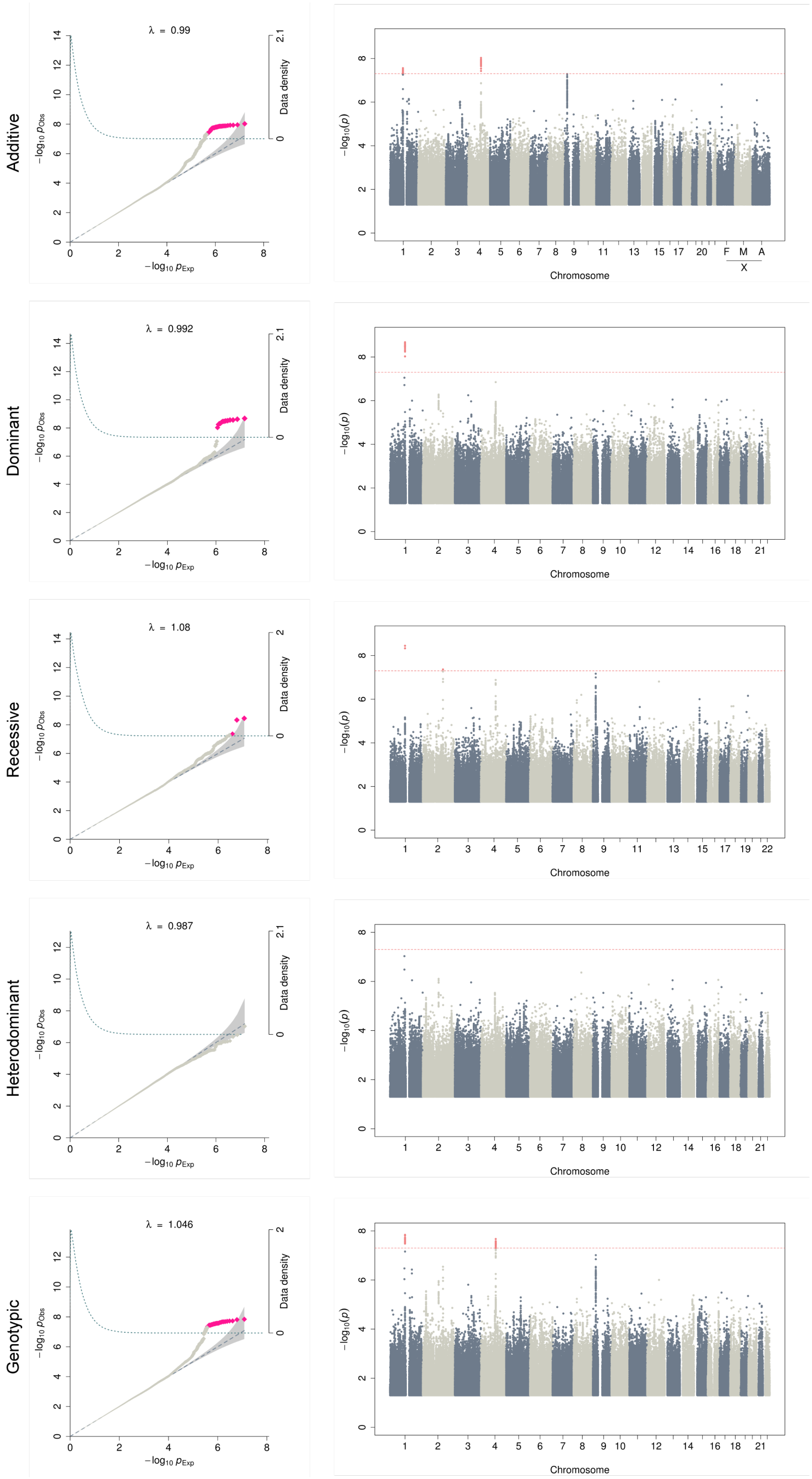


**Supplementary Figure 4. Q-Q plots and Manhattan plots for cardiovascular disease.**

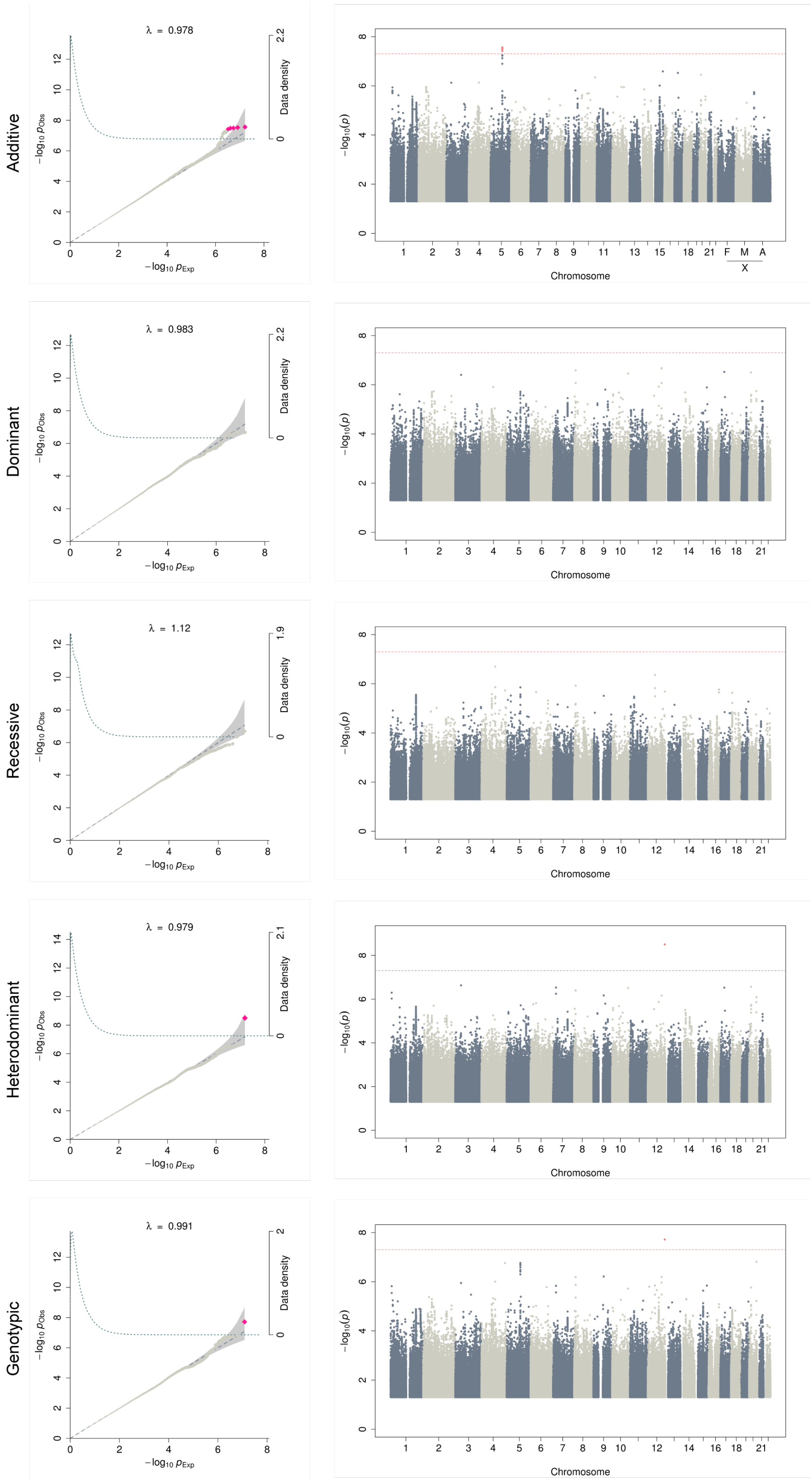


**Supplementary Figure 5. Q-Q plots and Manhattan plots for major depressive disorder.**

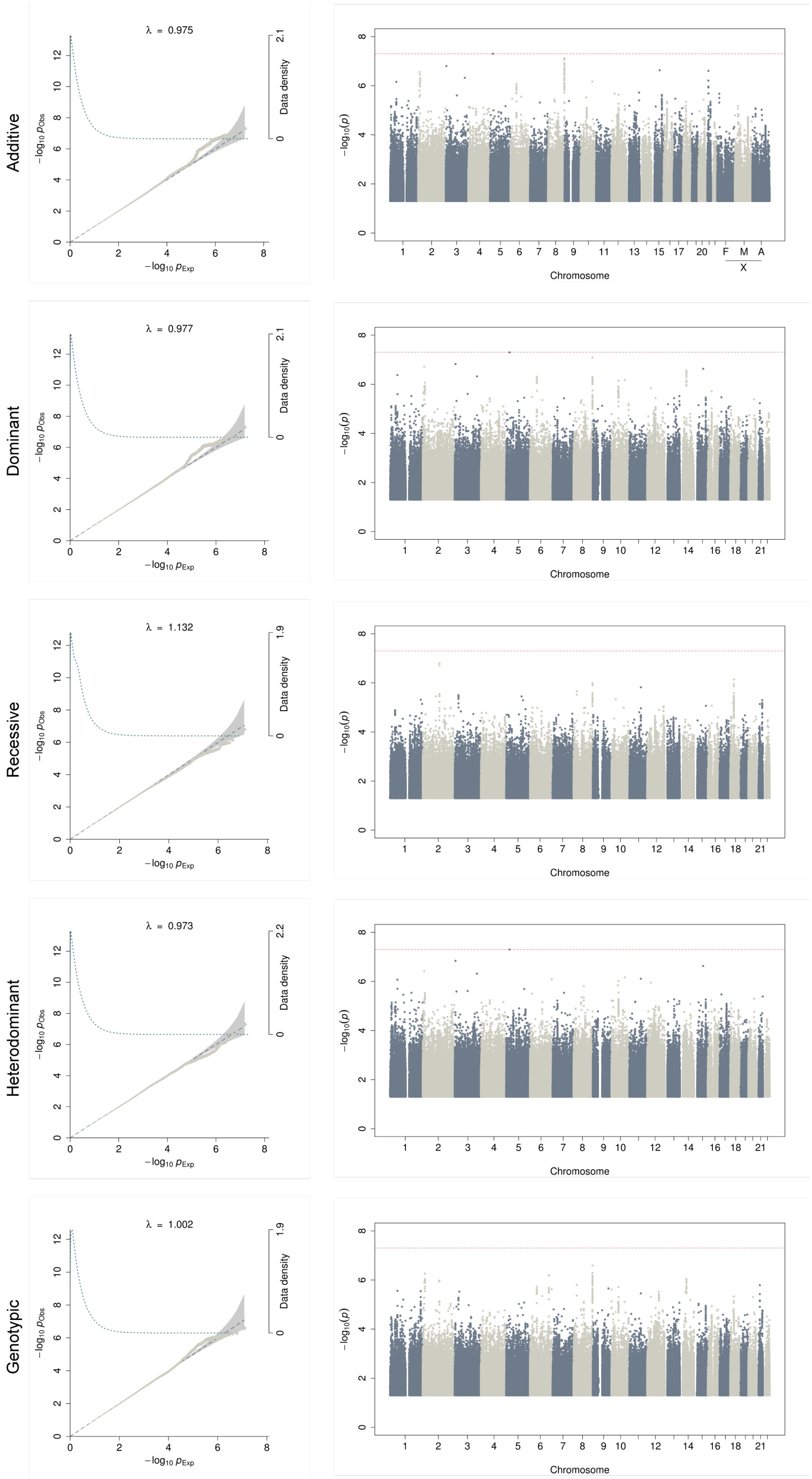


**Supplementary Figure 6. Q-Q plots and Manhattan plots for dermatophytosis.**

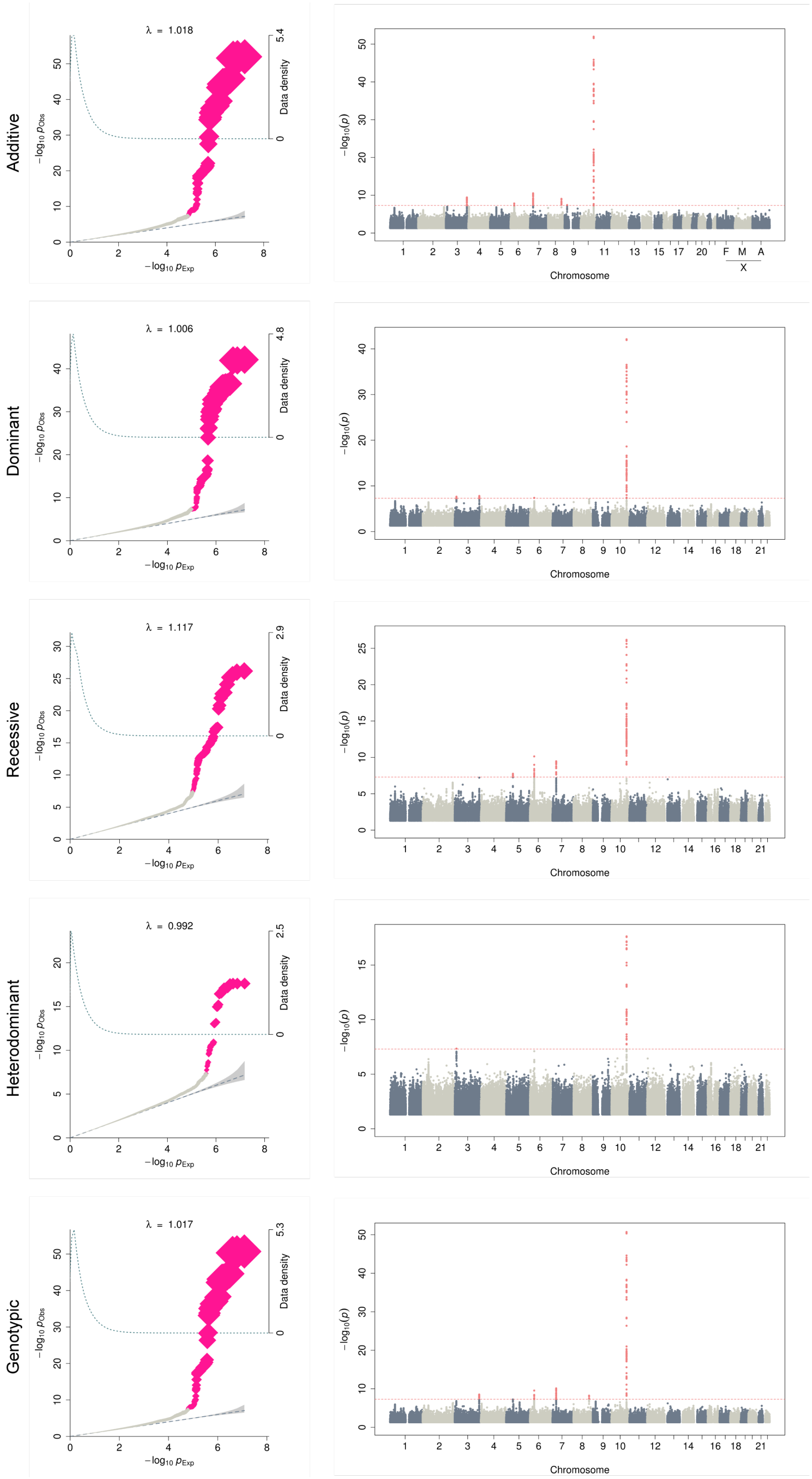


**Supplementary Figure 7. Q-Q plots and Manhattan plots for type 2 diabetes.**

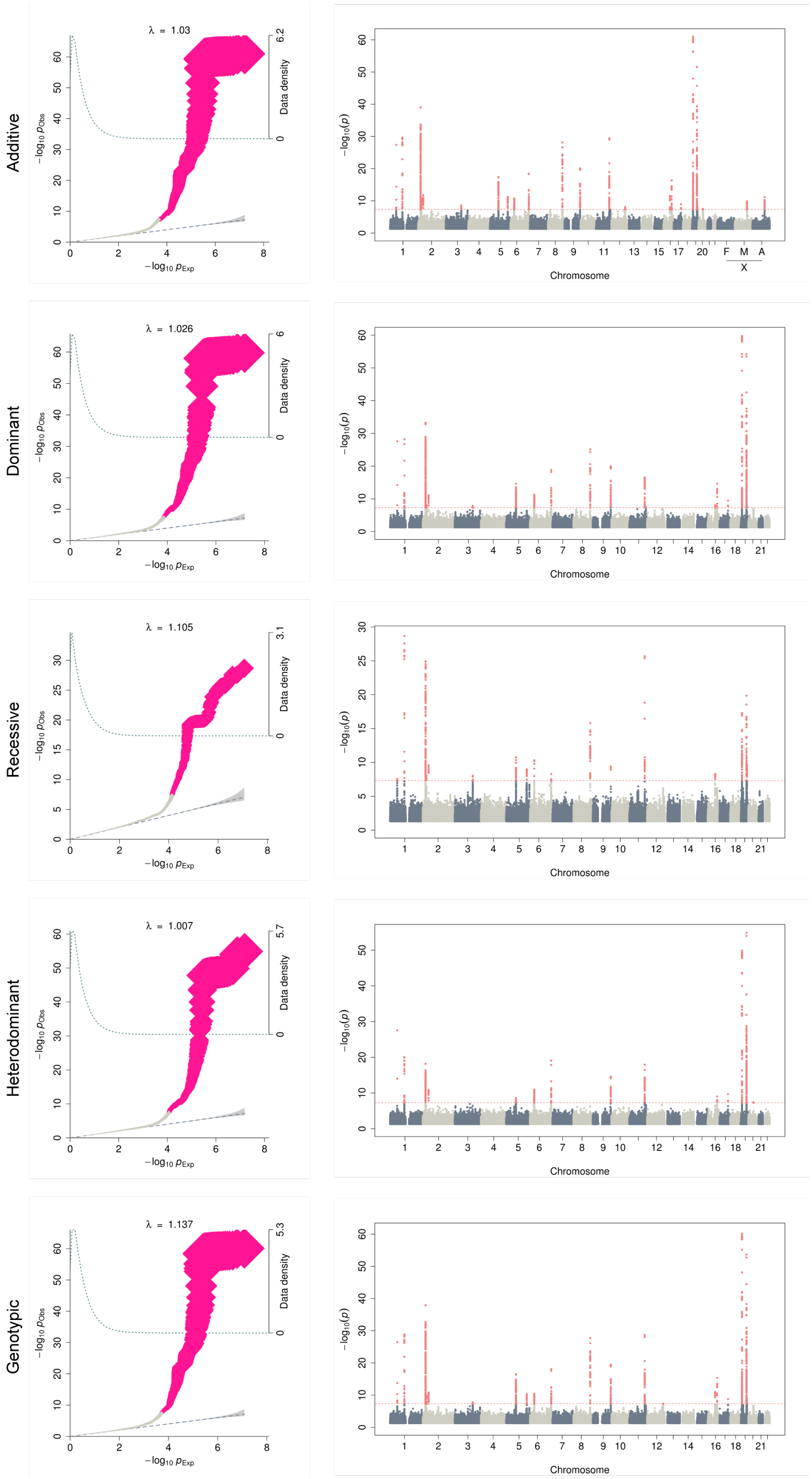


**Supplementary Figure 8. Q-Q plots and Manhattan plots for dyslipidemia.**

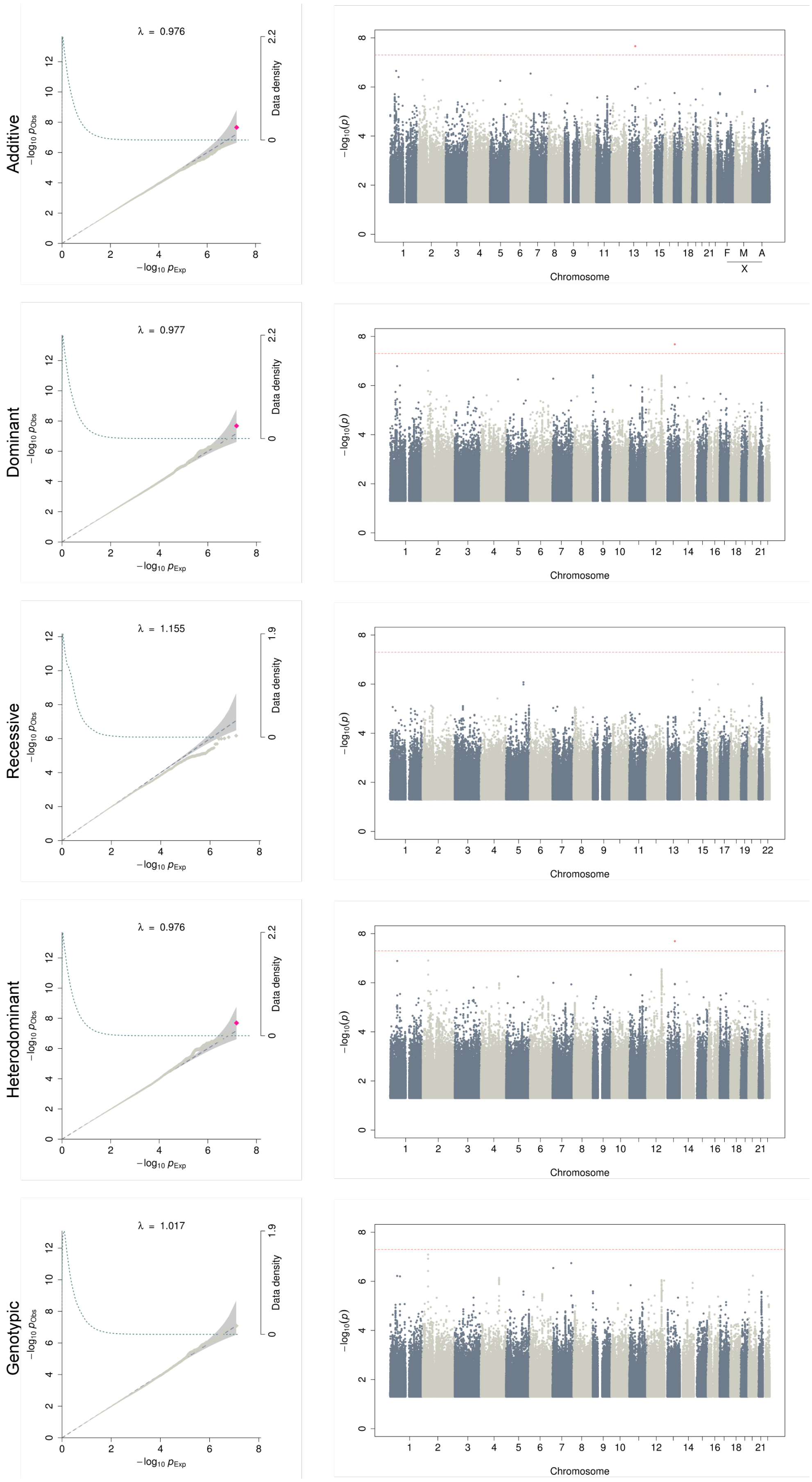


**Supplementary Figure 9. Q-Q plots and Manhattan plots for hemorrhoids.**

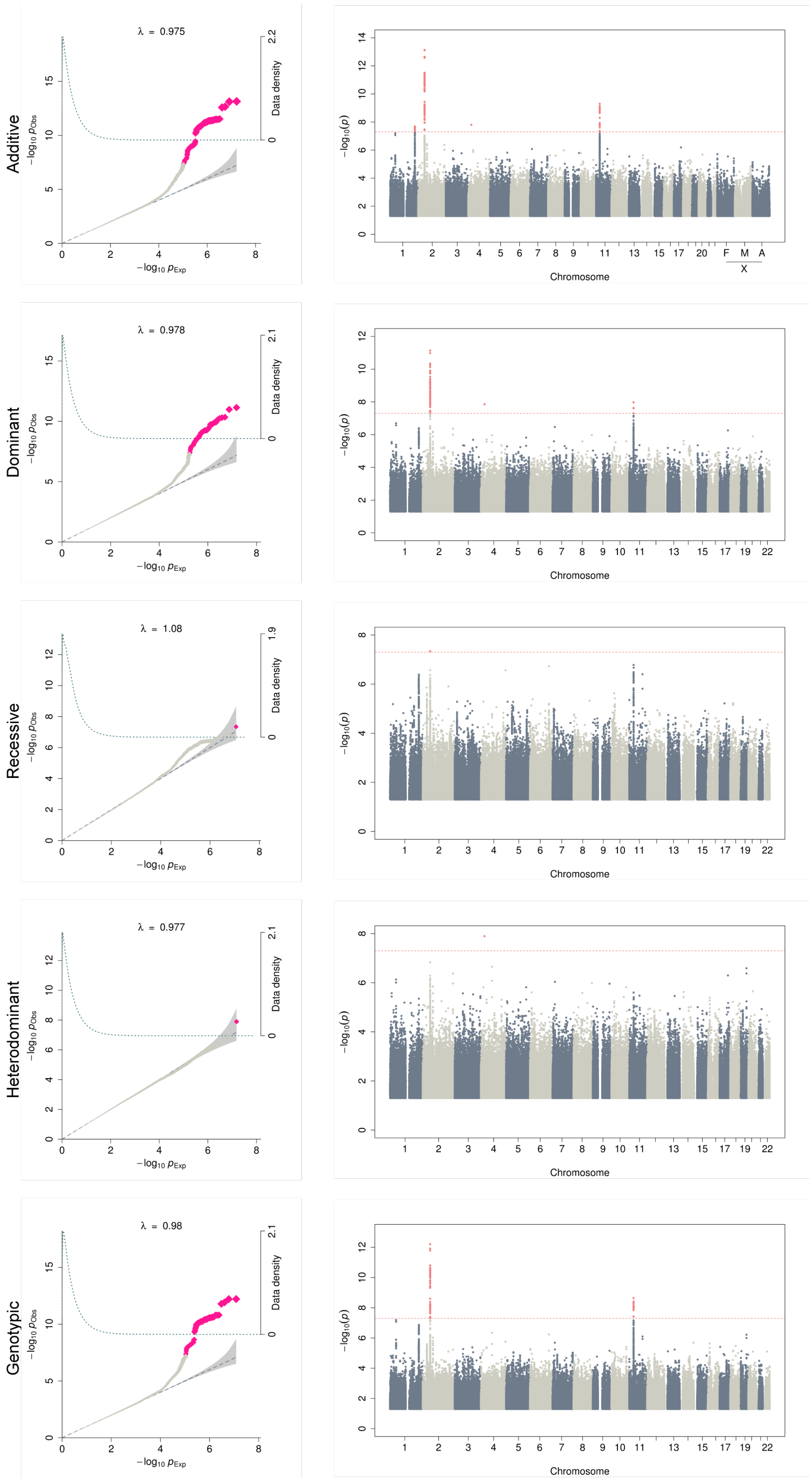


**Supplementary Figure 10. Q-Q plots and Manhattan plots for hernia abdominopelvic cavity.**

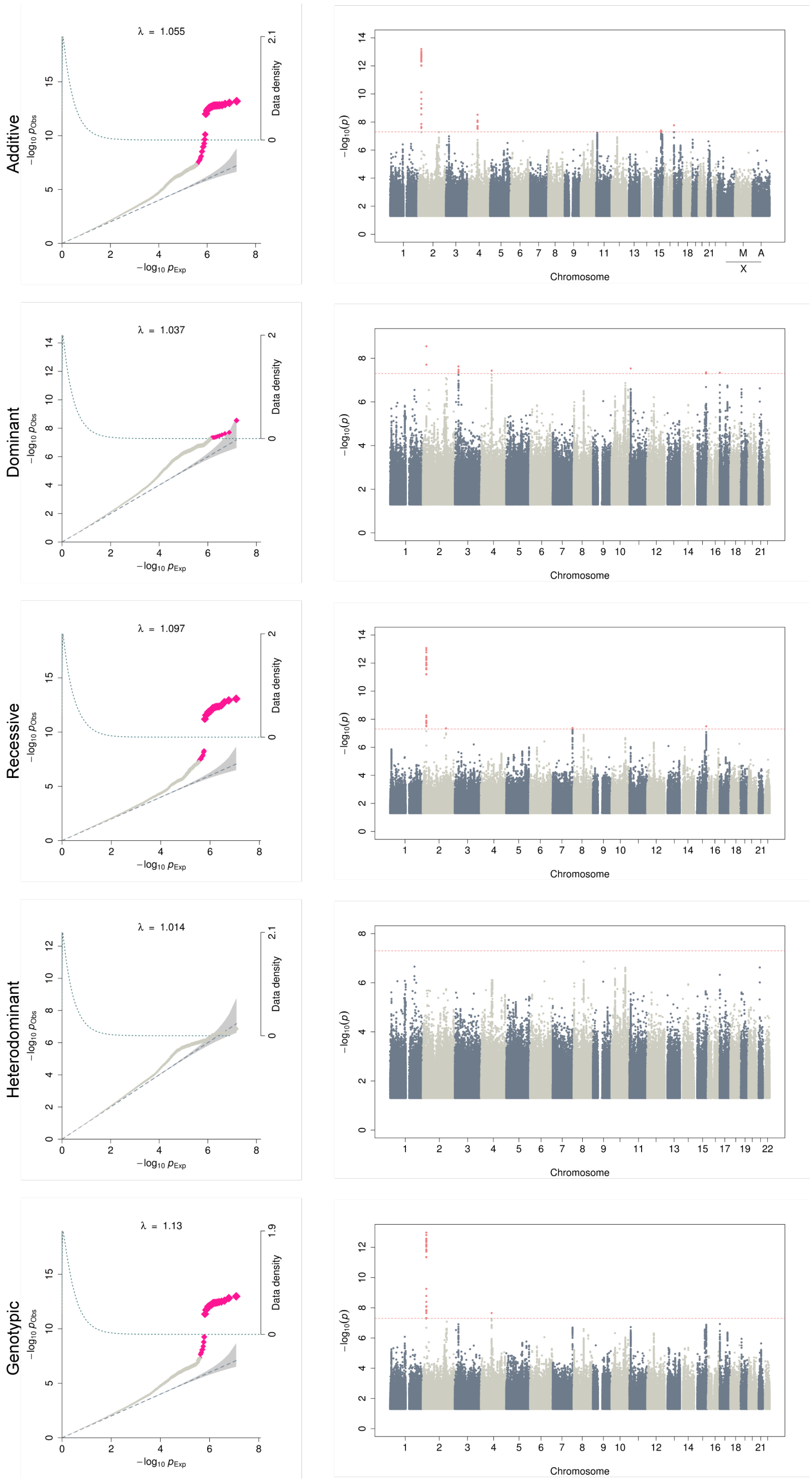


**Supplementary Figure 11. Q-Q plots and Manhattan plots for hypertension.**

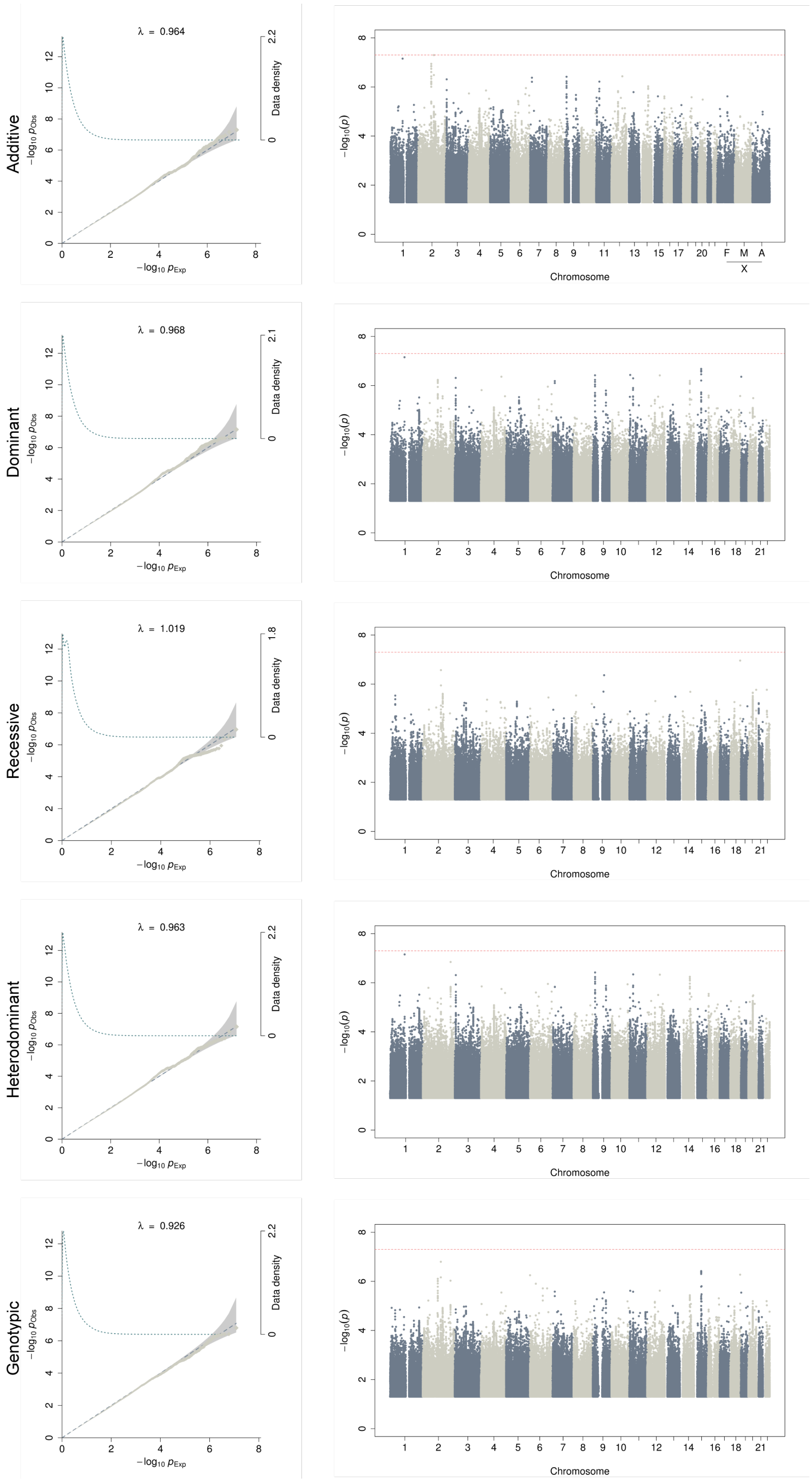


**Supplementary Figure 12. Q-Q plots and Manhattan plots for insomnia.**

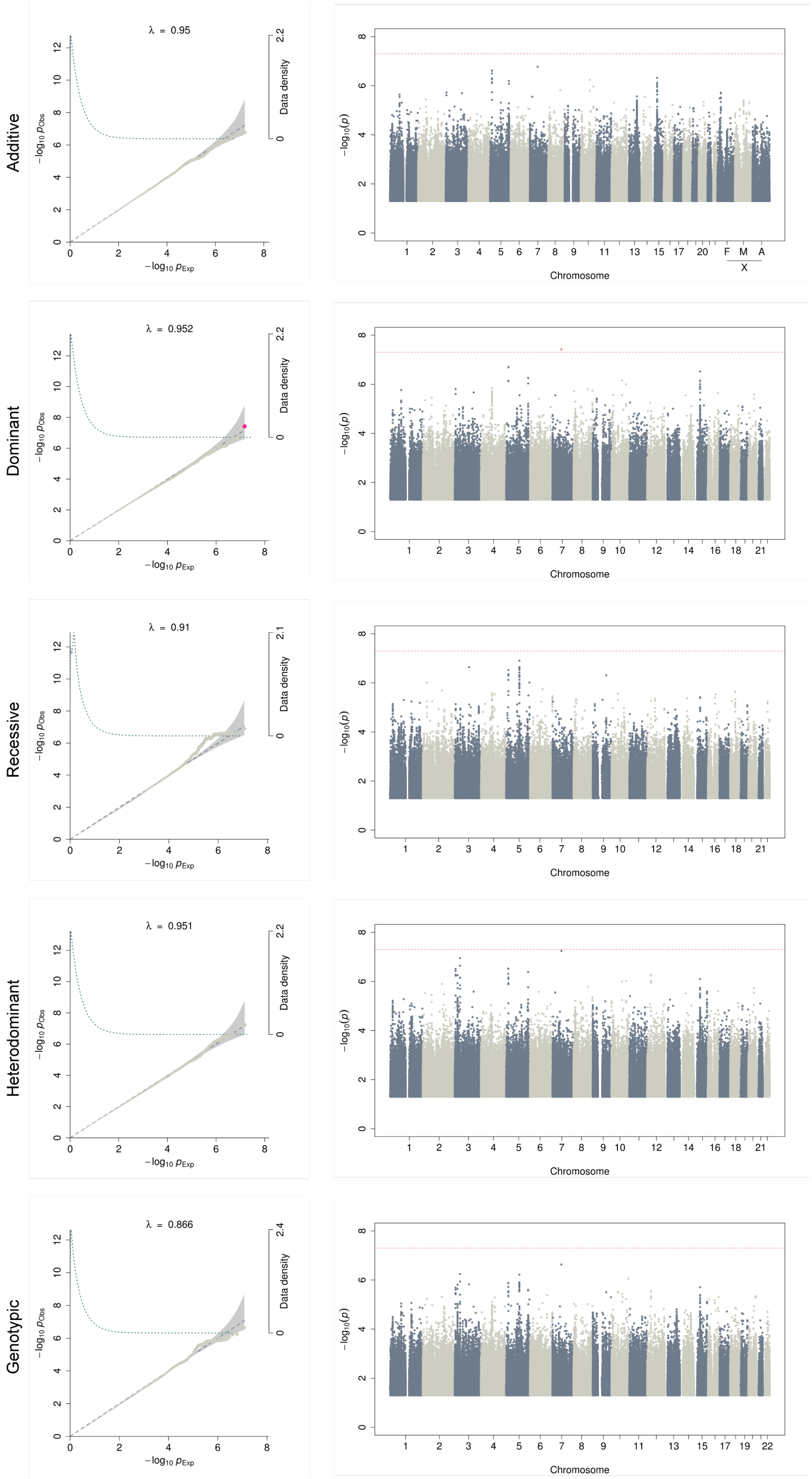


**Supplementary Figure 13. Q-Q plots and Manhattan plots for iron deficiency**

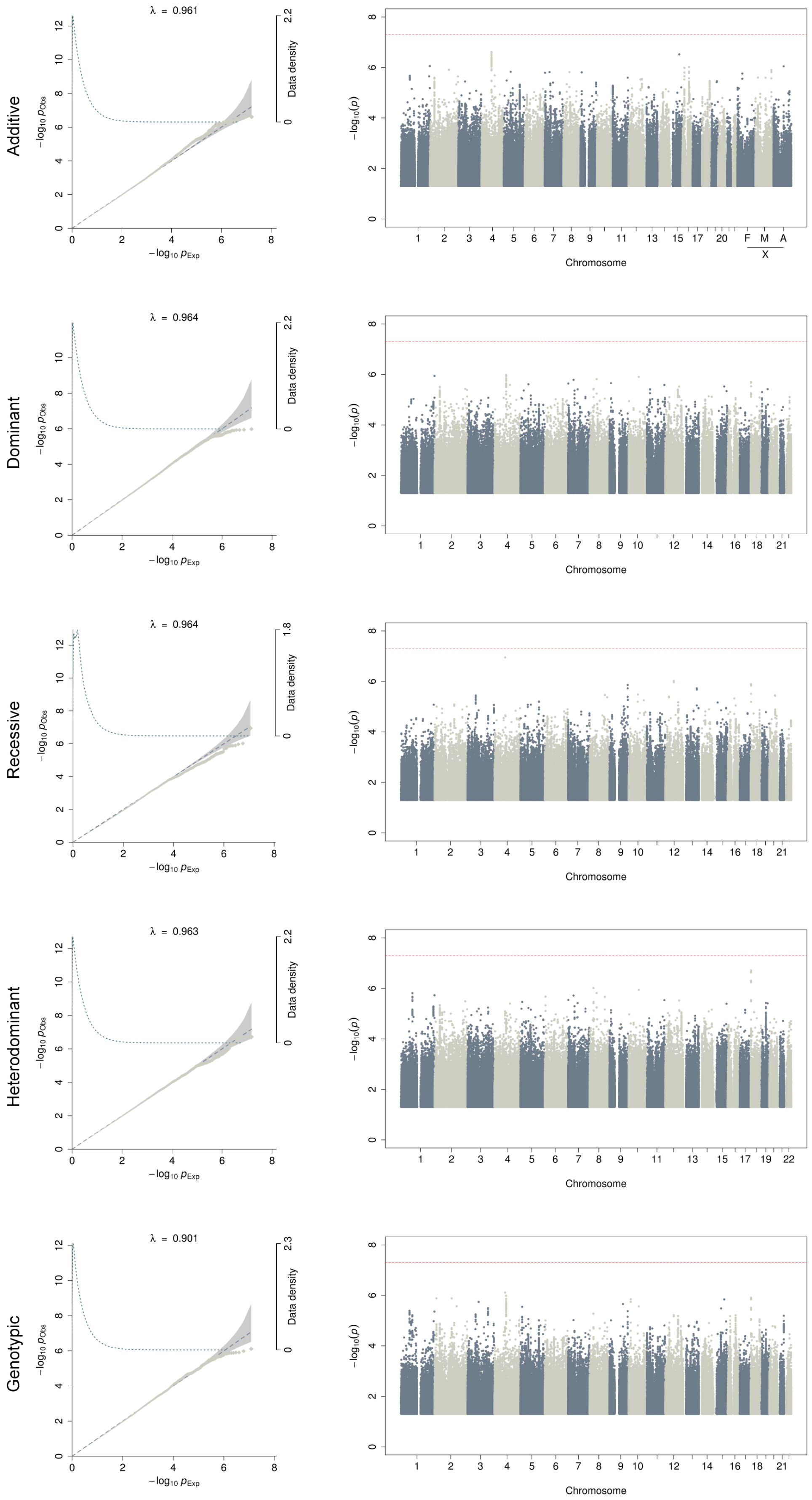


**Supplementary Figure 14. Q-Q plots and Manhattan plots for irritable bowel syndrome.**

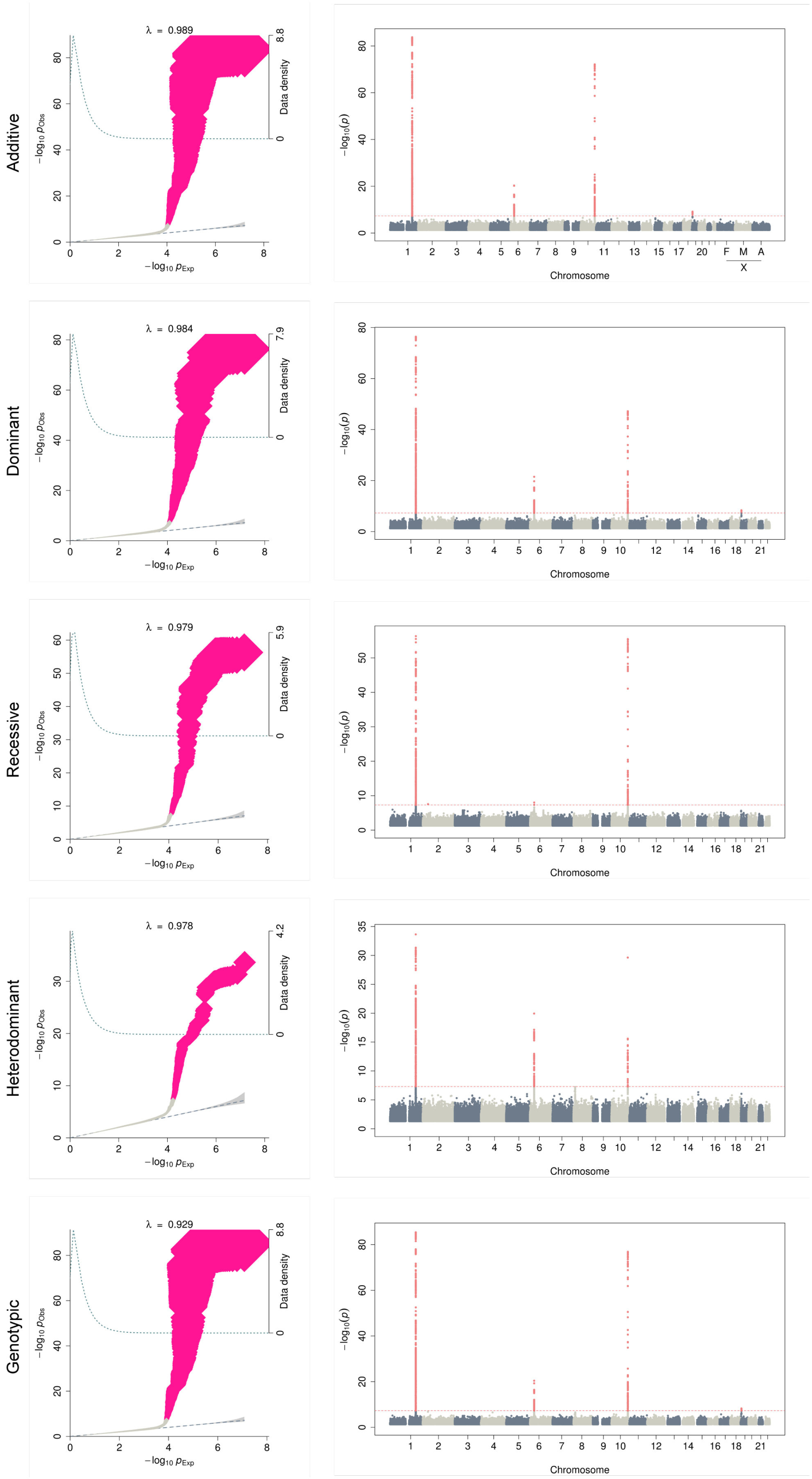


**Supplementary Figure 15. Q-Q plots and Manhattan plots for macular degeneration.**

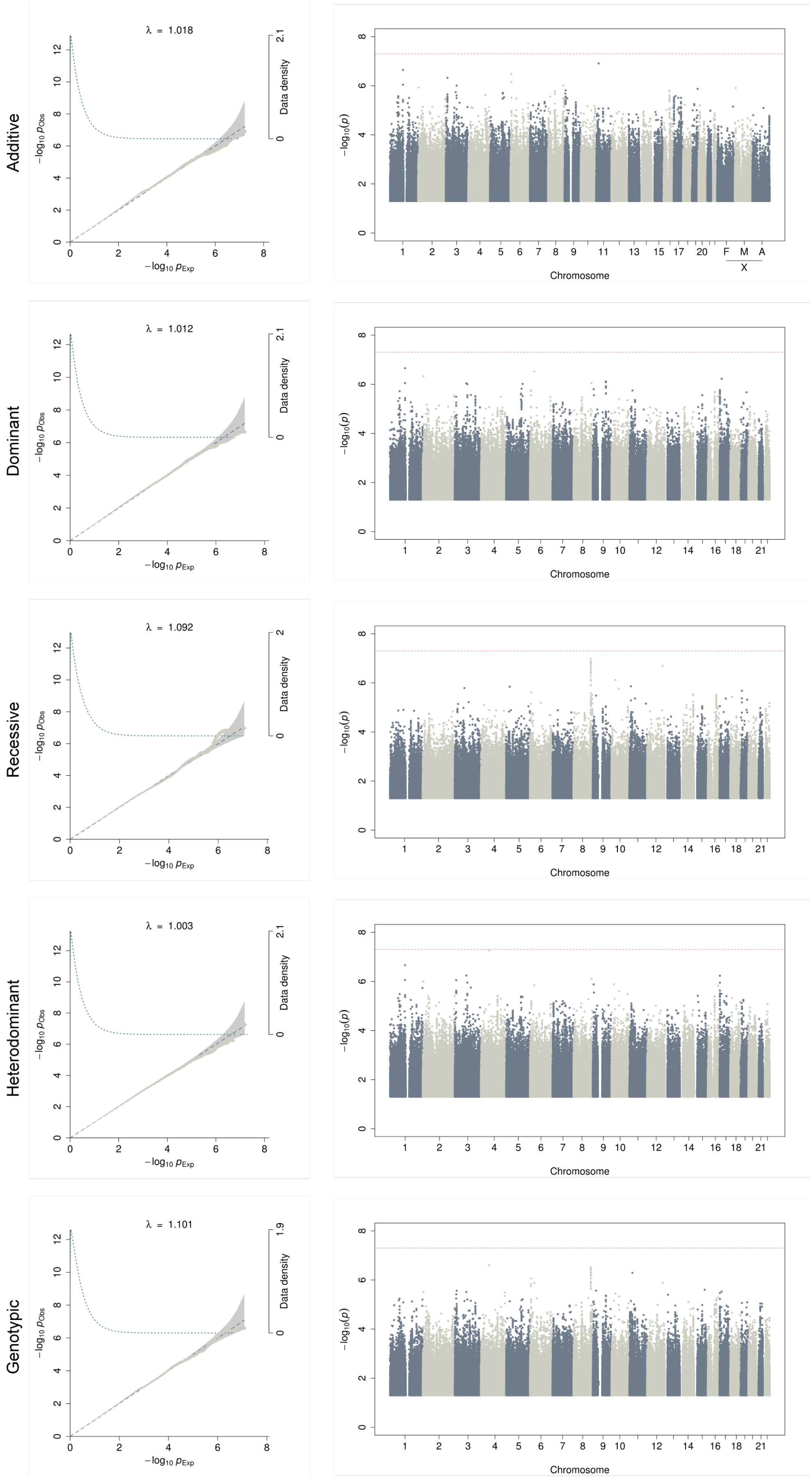


**Supplementary Figure 16. Q-Q plots and Manhattan plots for osteoarthritis.**

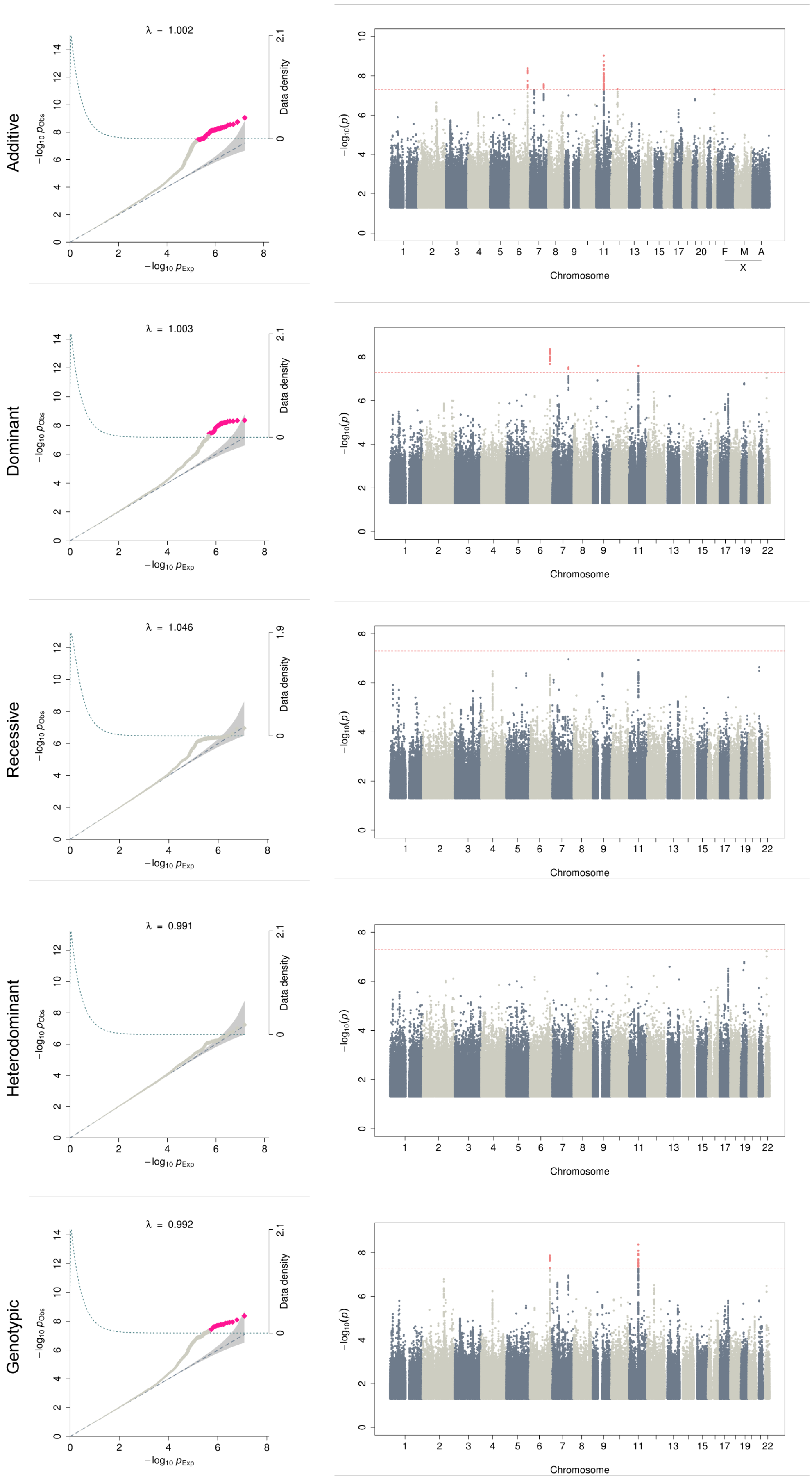


**Supplementary Figure 17. Q-Q plots and Manhattan plots for osteoporosis.**

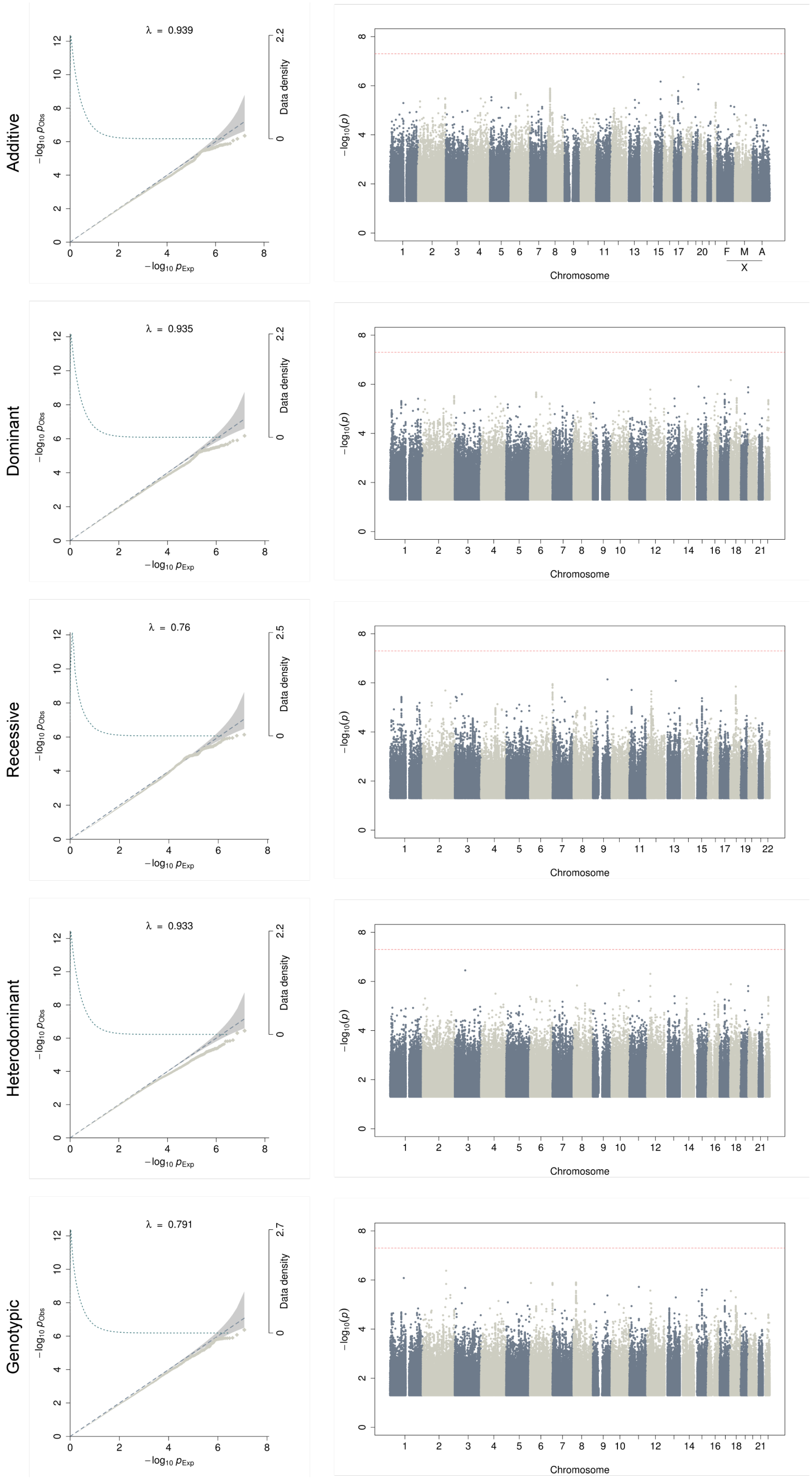


**Supplementary Figure 18. Q-Q plots and Manhattan plots for peptic ulcers.**

.

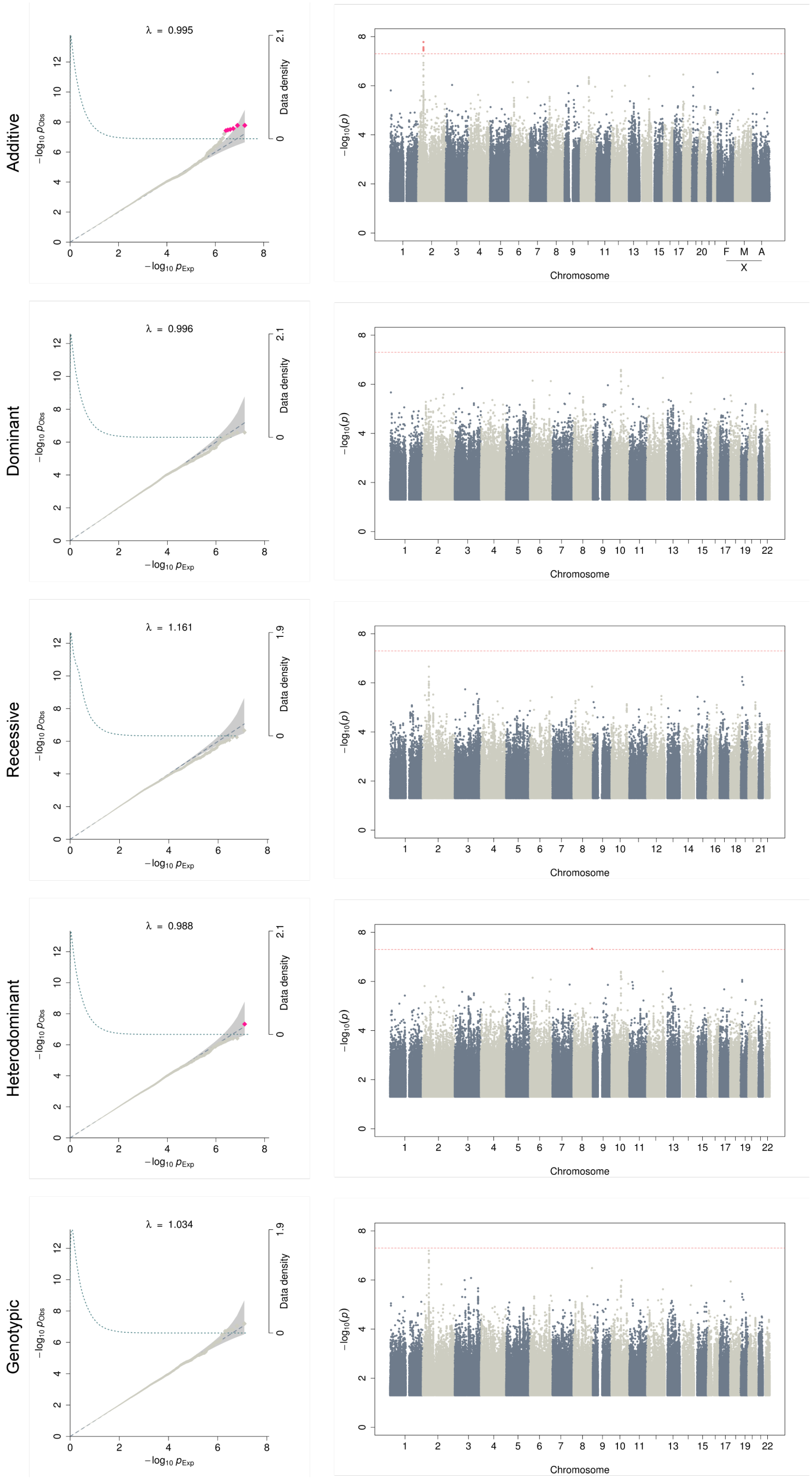


**Supplementary Figure 19. Q-Q plots and Manhattan plots for psychiatric disorders.**

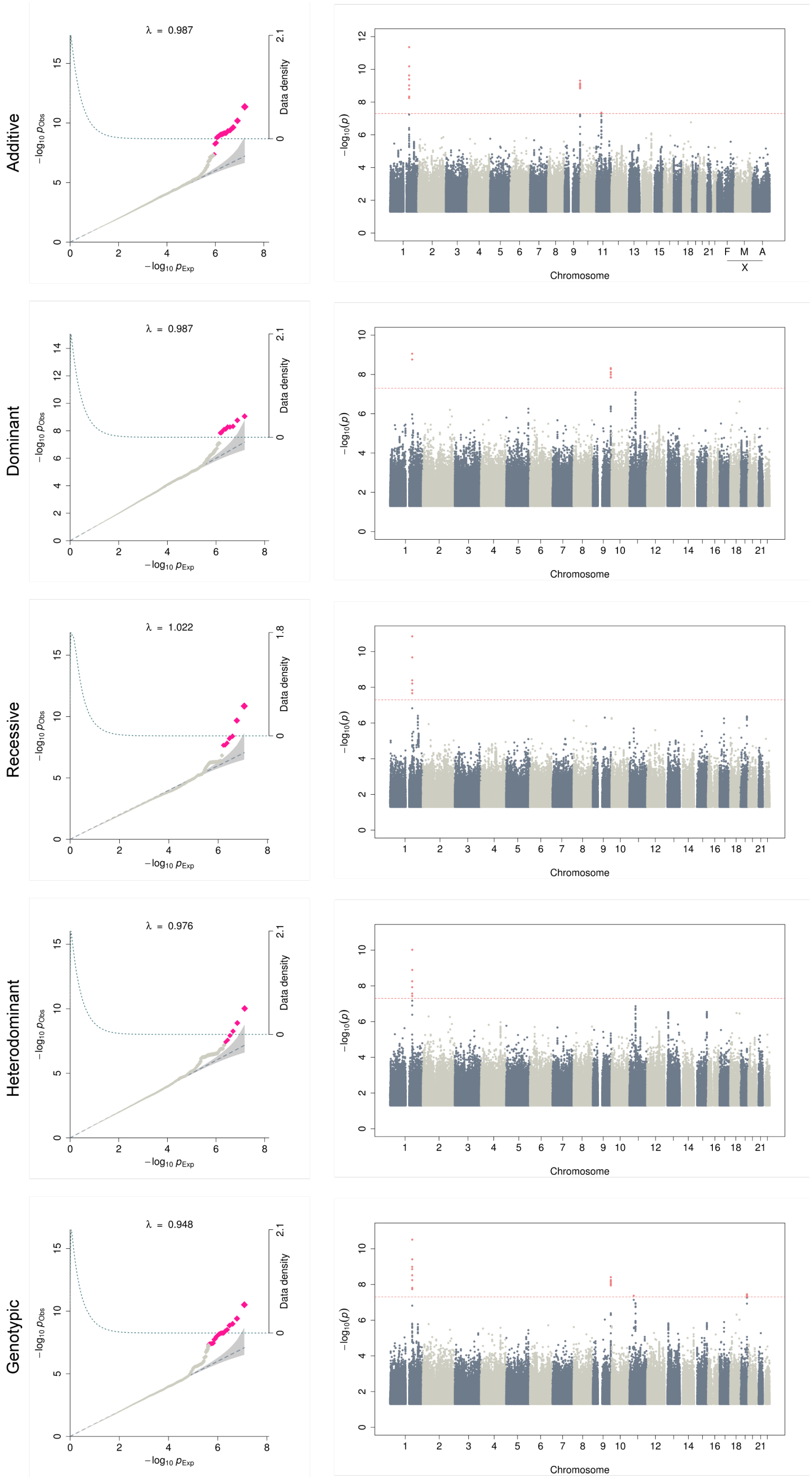


**Supplementary Figure 20. Q-Q plots and Manhattan plots for peripheral vascular disease.**

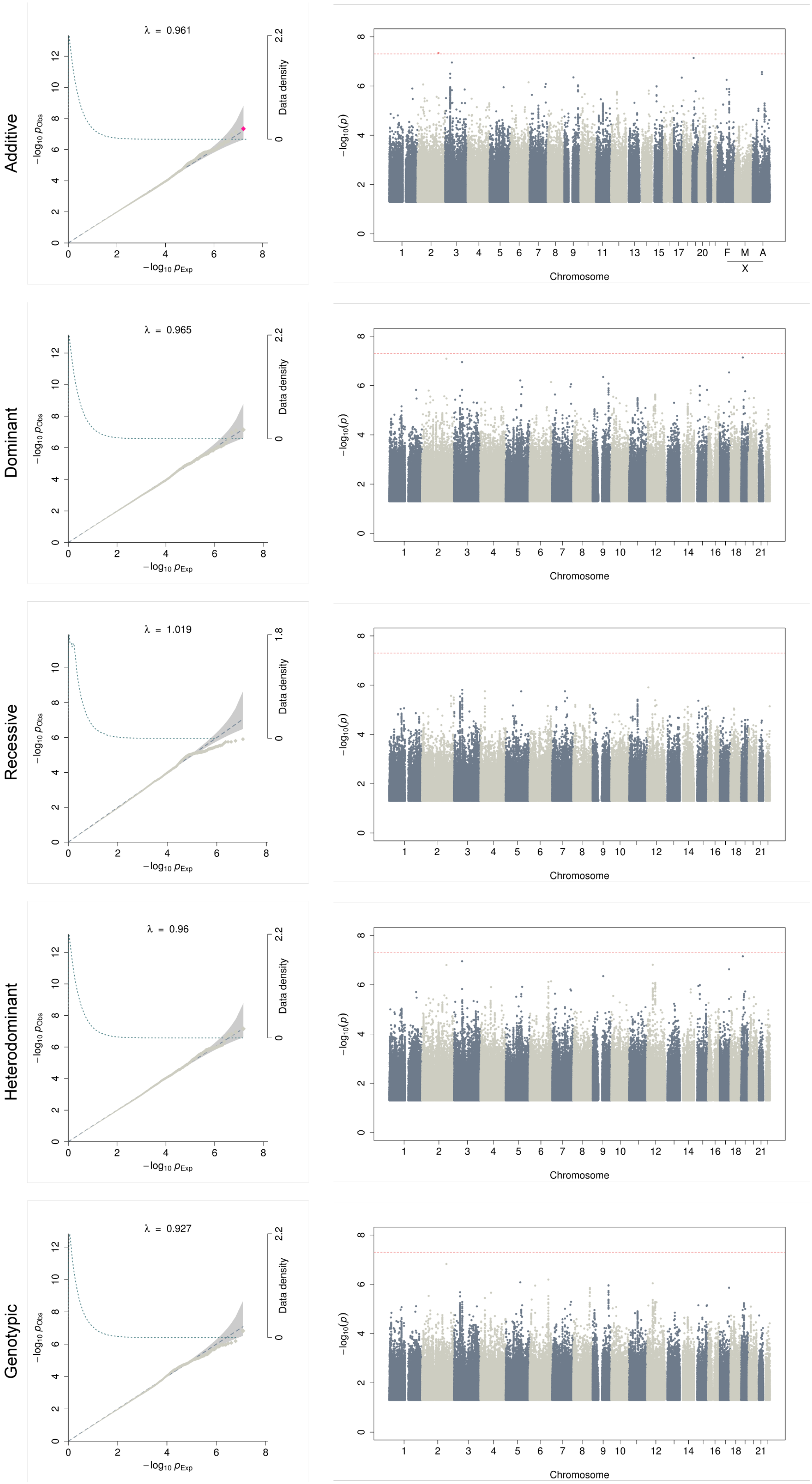


**Supplementary Figure 21. Q-Q plots and Manhattan plots for acute reaction to stress.**

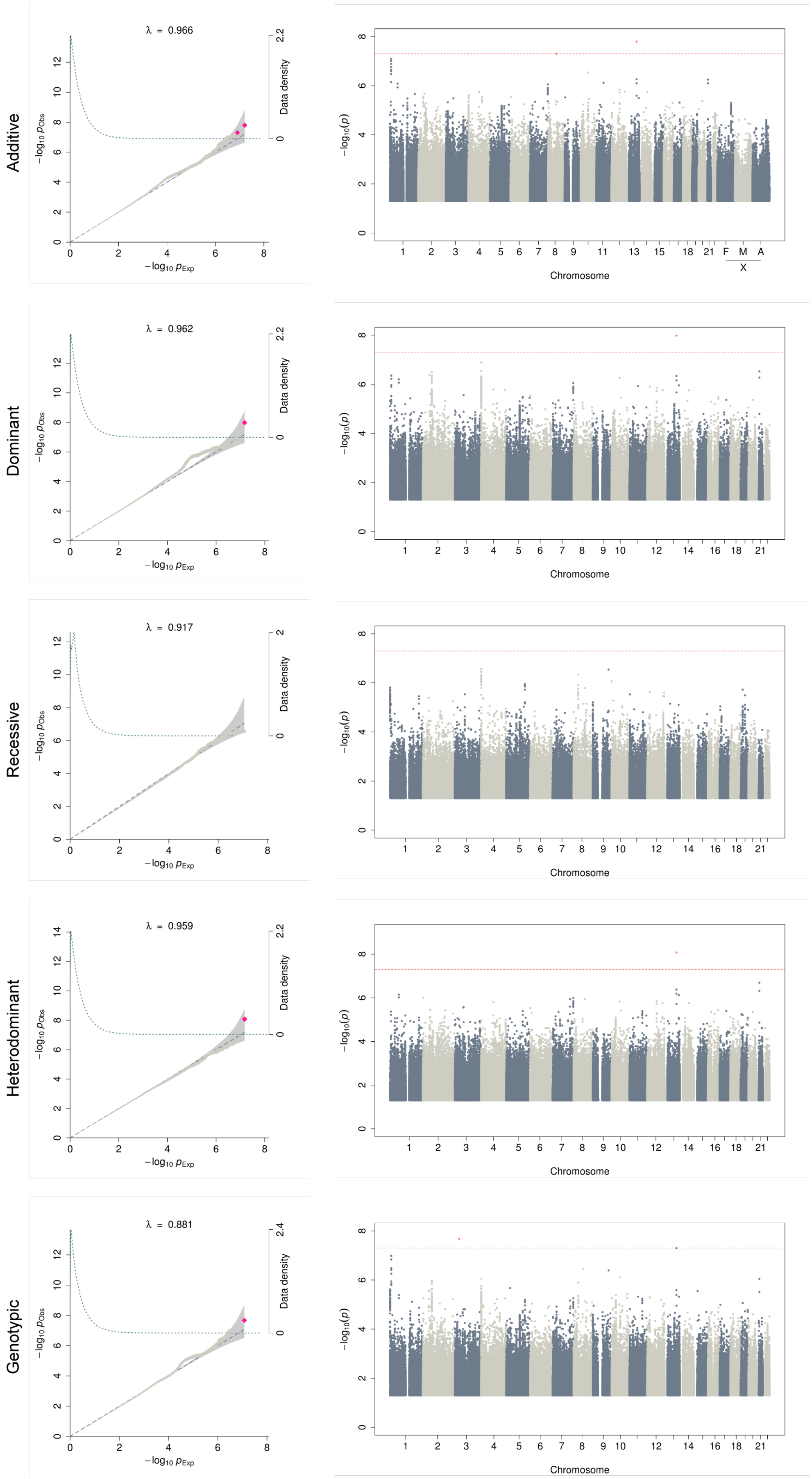


**Supplementary Figure 22. Q-Q plots and Manhattan plots for varicose veins.**
