## Supplementary figures and images for "The impact of non-additive genetic associations on age-related complex diseases"

### Supplementary Figure 23

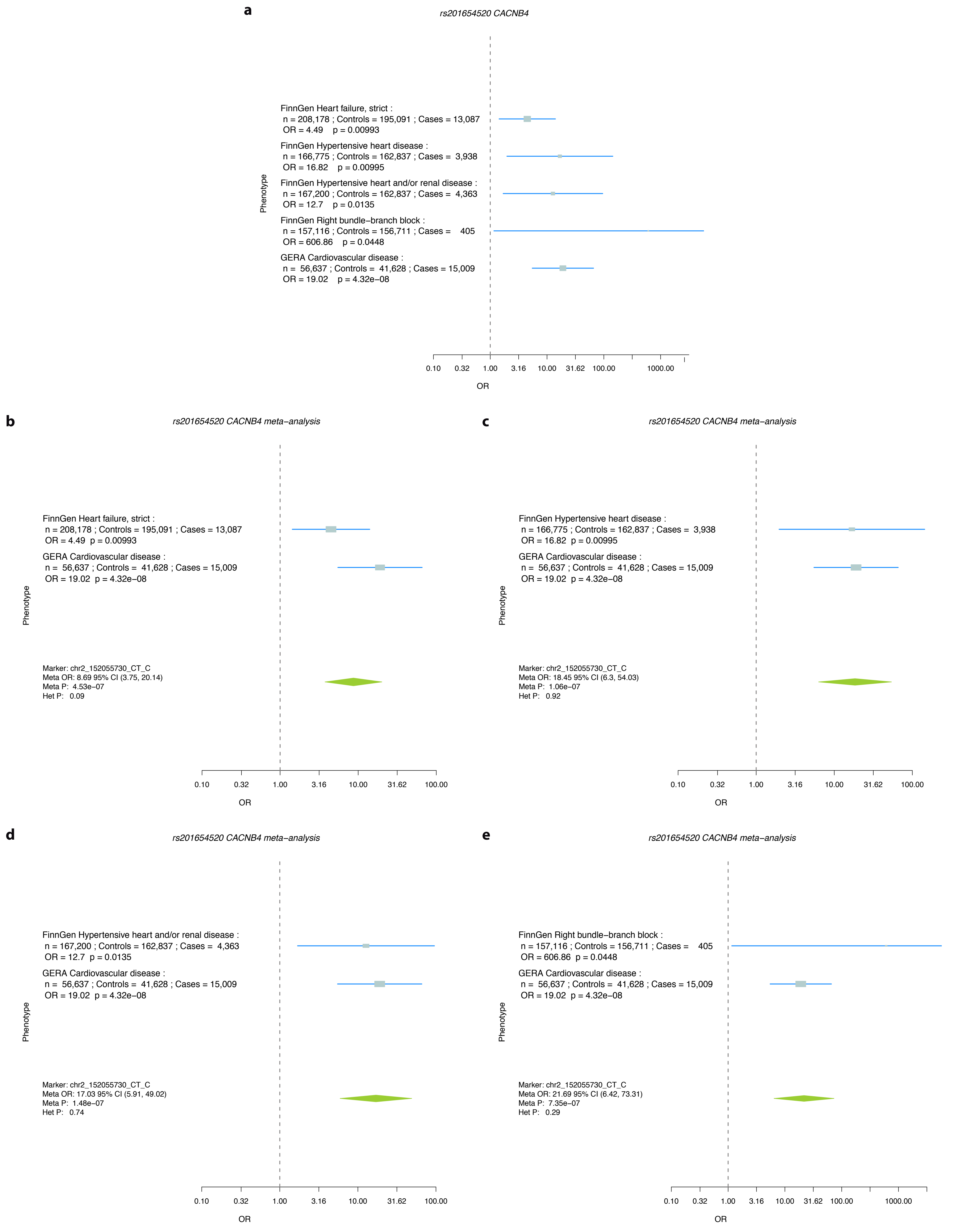

### Supplementary Figure 24

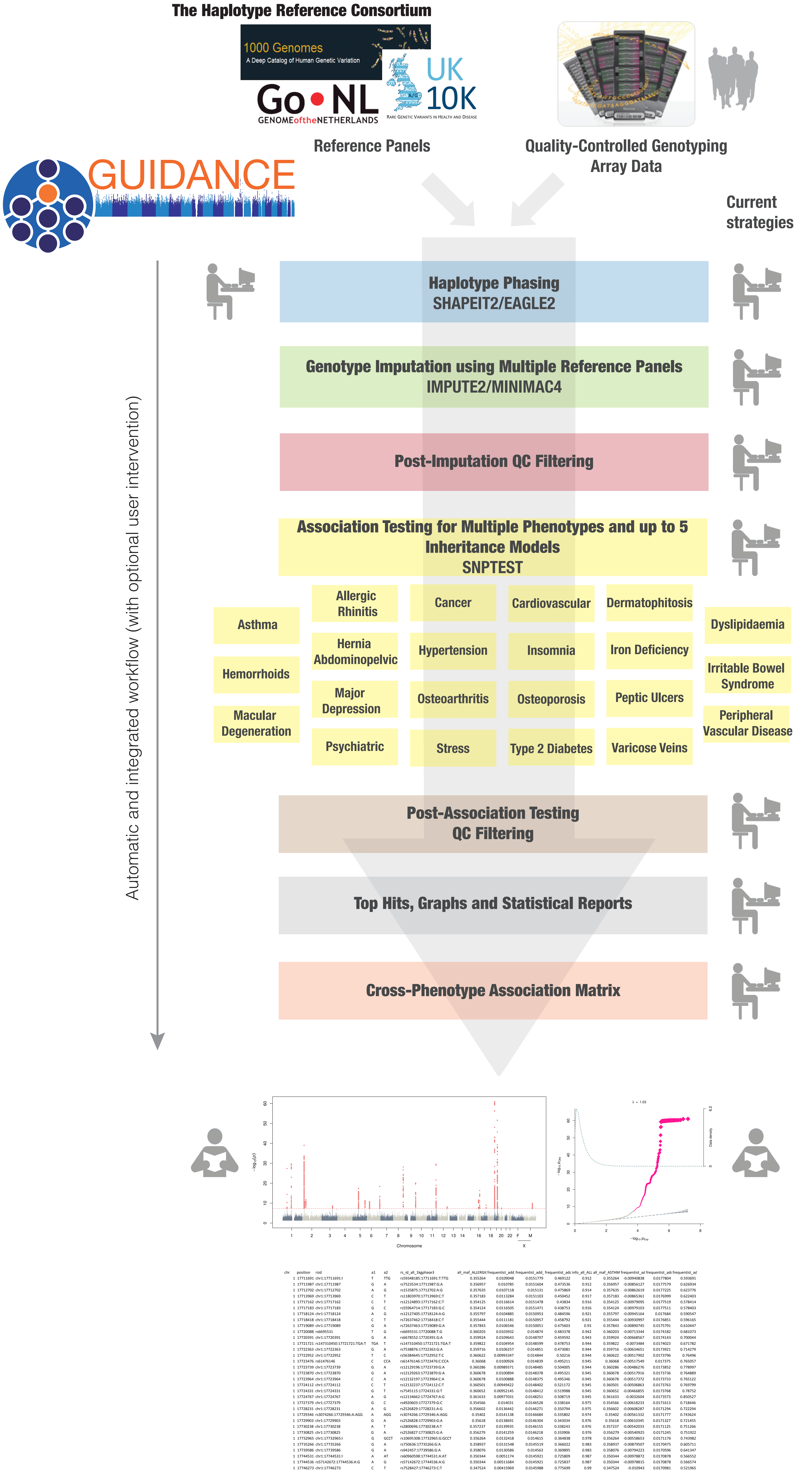

### Supplementary Figure 25

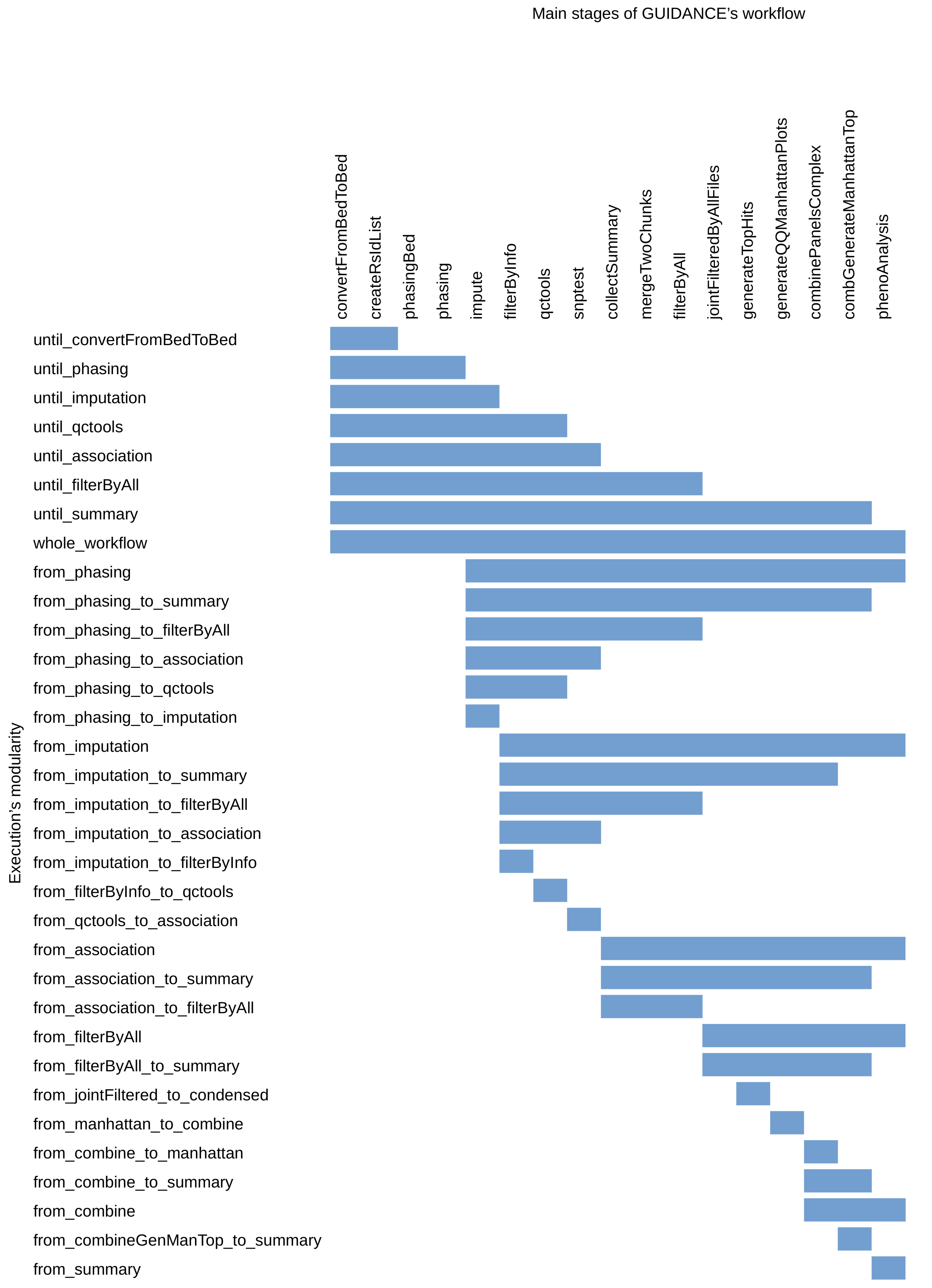

### Supplementary Figure 26

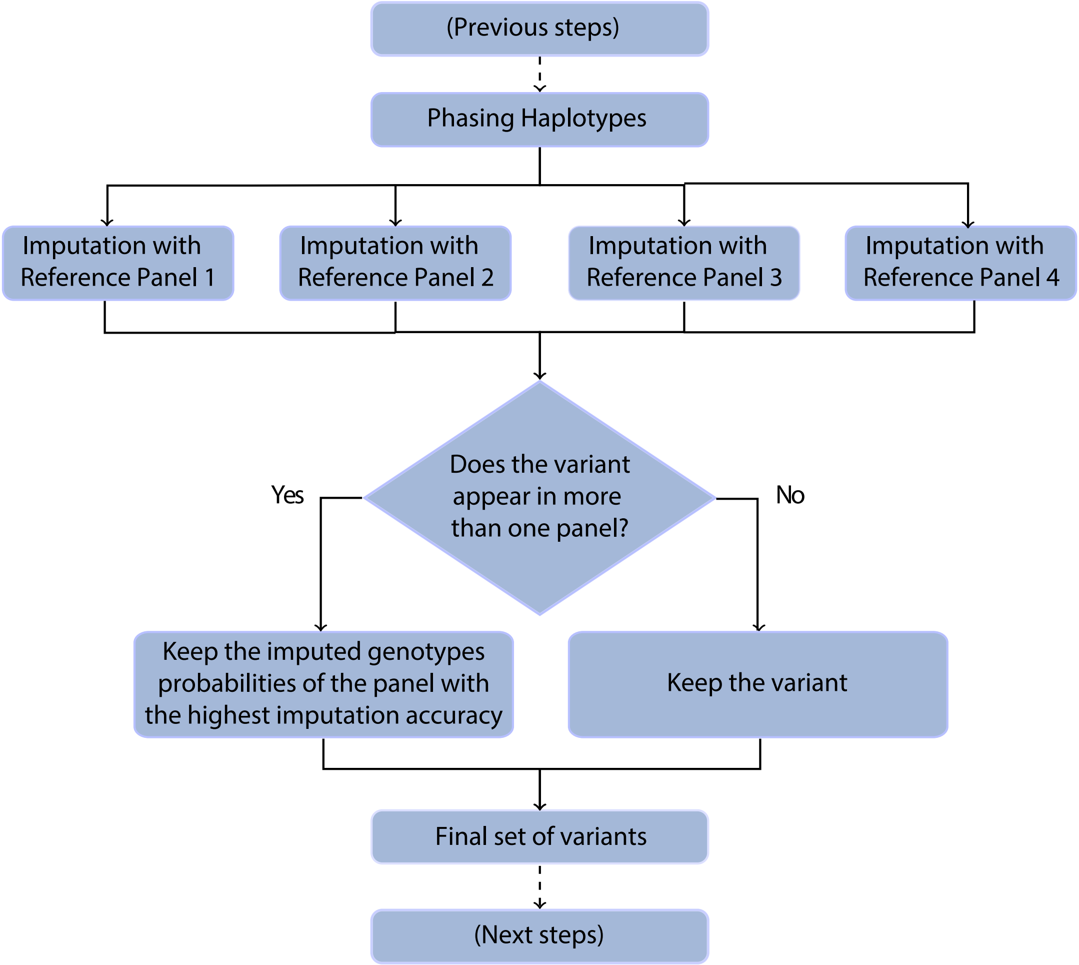

### Supplementary Figure 27

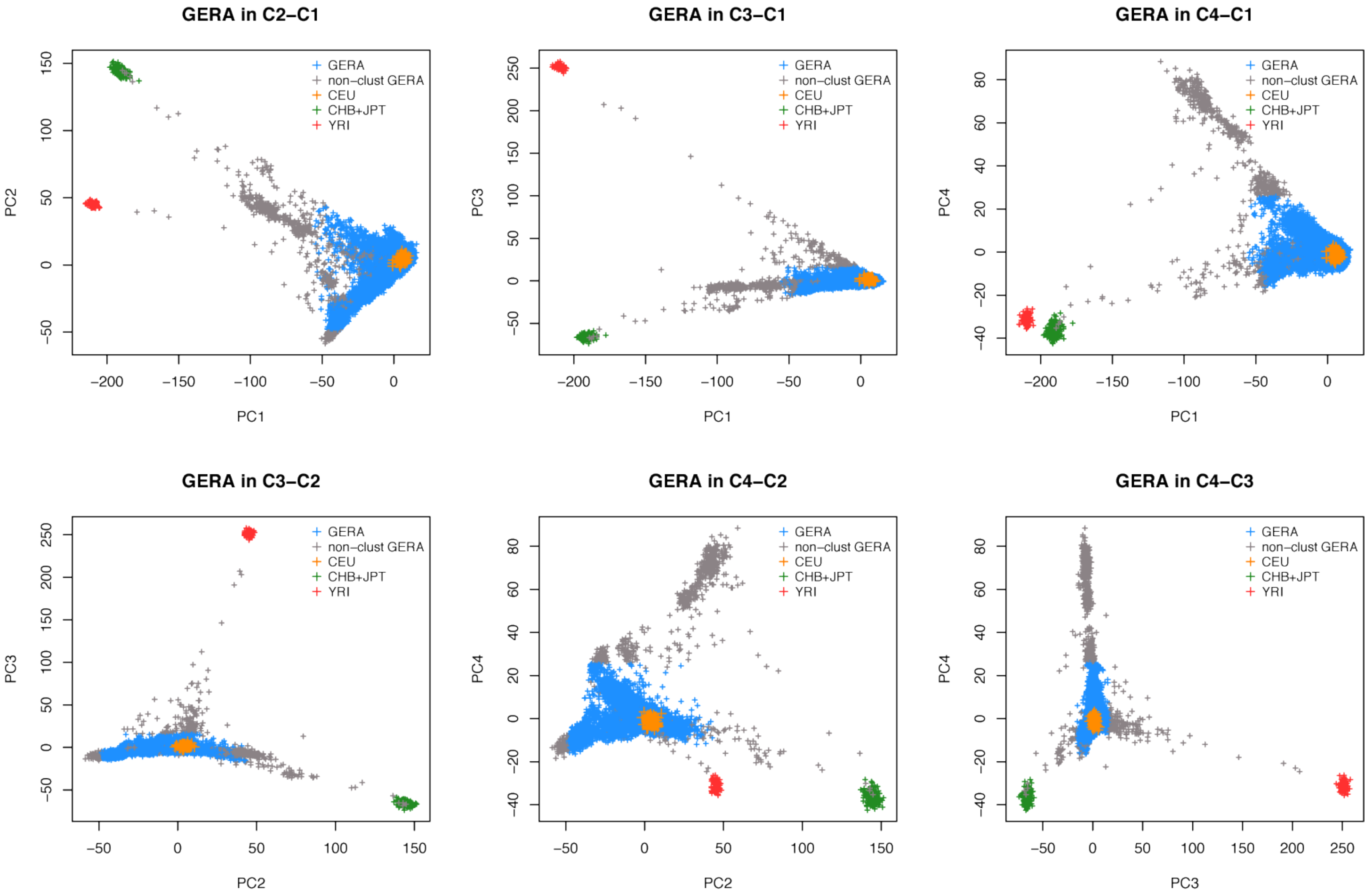

### Supplementary Figure 28

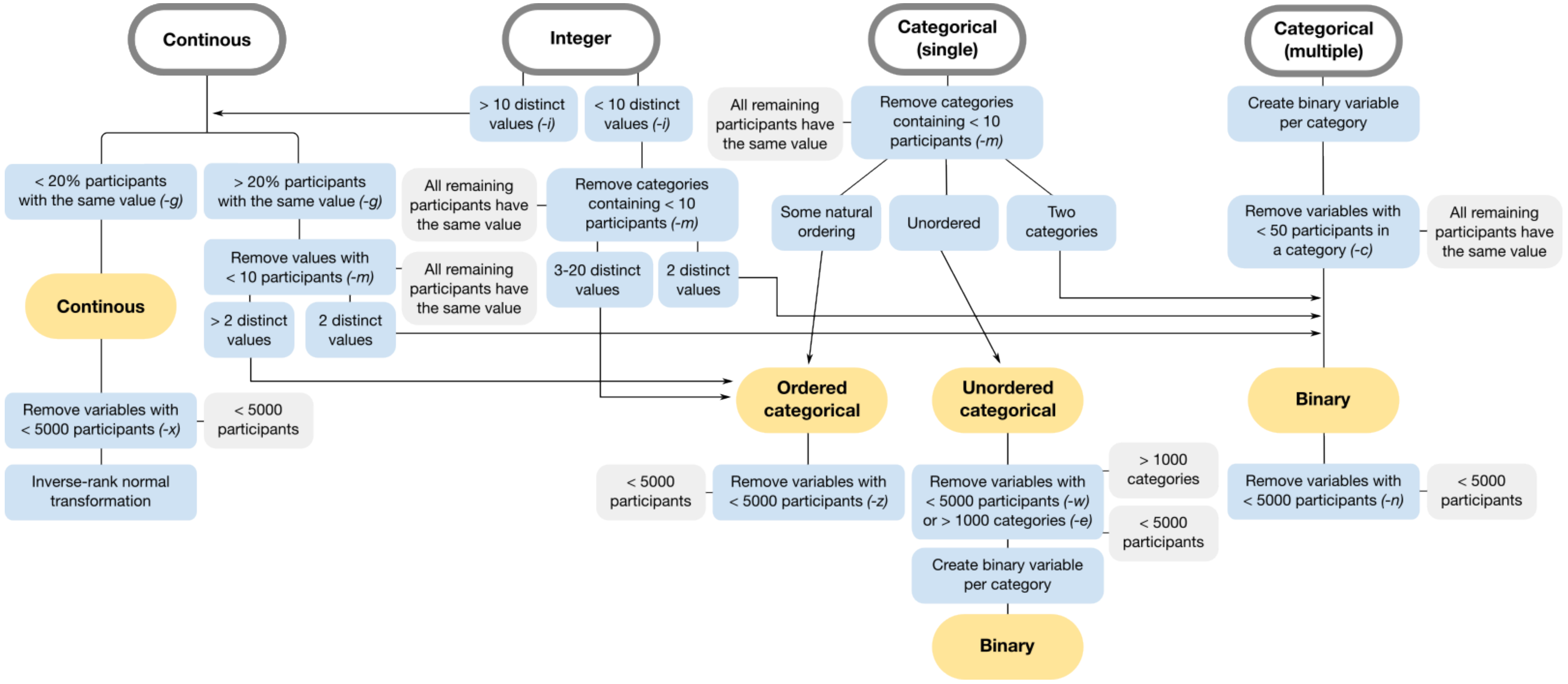
