## Supplementary Figure Legends for "The impact of non-additive genetic associations on age-related complex diseases"

**Supplementary Figure 1-22. QQ-plots and Manhattan plots from the 22 diseases analyzed after combining the results from 1000G phase 3, HRC, GoNL and UK10K, for additive, dominante, recessive, heterodominant and genotypic or general inheritance model**. At left, the QQ-plot represents the expected −log10 *p*-values under the null hypothesis in the x axis, while observed −log10 p-values are represented in the y axis. At right, the Manhattan plots, represents the −log10 p-values in the y axis and the chromosome position in the x axis.

**Supplementary Figure 23. Replication with FinnGen of the recessive rs201654520 indel in *CACNB4* associated with cardiovascular disease. a** Effect size and 95% confidence intervals for heart failure, hypertensive heart disease, hypertensive heart and/or renal disease, right bundle-branch block, and the results from the analysis of the GERA cohort. All the phenotypes show a direction of effect consistent with the effect observed in the GERA analysis. **b** Forest plot showing the meta-analysis of the association of rs201654520 with cardiovascular disease observed in the GERA cohort with hypertensive heart failure in FinnGen. **c** Forest plot showing the meta-analysis of the association of rs201654520 with cardiovascular disease observed in the GERA cohort with hypertensive heart disease in FinnGen. **d** Forest plot showing the meta-analysis of the association of rs201654520 with cardiovascular disease observed in the GERA cohort with hypertensive heart and/or renal disease in FinnGen. **e** Forest plot showing the meta-analysis of the association of rs201654520 with cardiovascular disease observed in the GERA cohort with right bundle-branch block in FinnGen.

**Supplementary Figure 24. Schematic representation of the steps included GUIDANCE compared to current GWAS workflows.** At both sides of the workflow, the steps that typically require manual intervention in current strategies are displayed (right) and compared with GUIDANCE requirements of user intervention (left), which allows an automatic execution. GUIDANCE starts with Quality Controlled genetic data (top), following the haplotypes phasing, genotype imputation using multiple panels, and association testing considering multiple phenotypes and inheritance models. GUIDANCE finishes with summary statistics and graphical representation of the results (bottom). Multiple genotypes displayed correspond to those found in GERA.

**Supplementary Figure 25. Modularity of GUIDANCE workflow.** The user can choose between running the whole workflow or just a subset of stages in the configuration file. This figure show the steps executed for each one of the acronyms that can be specified in the configuration file of GUIDANCE. The bar represents the stages (top) that will be executed by each category (left).

**Supplementary Figure 26. Flow-chart representation of how the results from several panels are combined to generate integrated results**. In case that one variant is present in more than one panel, the genotype probabilites that result from imputation with the best IMPUTE2 info score are selected. If a given variant is only imputed in one panel, the genotypes are selected from that panel. Only variants with an IMPUTE2 info score higher than the threshold that is specified in the configuration file (i. e., 0.7) are included in the final output.

**Supplementary Figure 27. The first 4 PCs derived from the GERA cohort superimposed on top of HapMap3 individuals.** Individuals that do not cluster with the European samples from HapMap are depicted in grey and removed from the dataset. Samples in blue represent the GERA cohort individuals of European self-reported ancestry that remain for further analysis.

**Supplementary Figure 28. Phenotype curation pipeline.** Raw phenotype data (gray outlined boxes) are passed to PHESANT, and a collection of filters (blue boxes) are applied. The thresholds shown here are the defaults in our modified version of PHESANT that can be altered in the phenomescan.r code using the flags displayed in parentheses. Gray filled boxes display the criteria for removal, and yellow filled boxes show the category of the variable after the rules in the blue boxes have been enforced.
